# A dynamic generative model can extract interpretable oscillatory components from multichannel neurophysiological recordings

**DOI:** 10.1101/2023.07.26.550594

**Authors:** Proloy Das, Mingjian He, Patrick L. Purdon

## Abstract

Modern neurophysiological recordings are performed using multichannel sensor arrays that are able to record activity in an increasingly high number of channels numbering in the 100’s to 1000’s. Often, underlying lower-dimensional patterns of activity are responsible for the observed dynamics, but these representations are difficult to reliably identify using existing methods that attempt to summarize multivariate relationships in a post-hoc manner from univariate analyses, or using current blind source separation methods. While such methods can reveal appealing patterns of activity, determining the number of components to include, assessing their statistical significance, and interpreting them requires extensive manual intervention and subjective judgement in practice. These difficulties with component selection and interpretation occur in large part because these methods lack a generative model for the underlying spatio-temporal dynamics. Here we describe a novel component analysis method anchored by a generative model where each source is described by a bio-physically inspired state space representation. The parameters governing this representation readily capture the oscillatory temporal dynamics of the components, so we refer to it as Oscillation Component Analysis (OCA). These parameters – the oscillatory properties, the component mixing weights at the sensors, and the number of oscillations – all are inferred in a data-driven fashion within a Bayesian framework employing an instance of the expectation maximization algorithm. We analyze high-dimensional electroencephalography and magnetoencephalography recordings from human studies to illustrate the potential utility of this method for neuroscience data.

**Significance Statement:** Neuroscience studies often involve simultaneous recordings in a large number of sensors in which a smaller number of dynamic components generate the complex spatio-temporal patterns observed in the data. Current blind source separation techniques produce sub-optimal results and are difficult to interpret because these methods lack an appropriate generative model that can guide both statistical inference and interpretation. Here we describe a novel component analysis method employing a dynamic generative model that can decompose high-dimensional multivariate data into a smaller set of oscillatory components are learned in a data-driven way, with parameters that are immediately interpretable. We show how this method can be applied to neurophysiological recordings with millisecond precision that exhibit oscillatory activity such as electroencephalography and magnetoencephalography.

## Introduction

Human neurophysiological recordings such as scalp electroencephalogram (EEG), magnetoencephalogram (MEG), stereoelectroencephalogram (SEEG), Local field potentials (LFP) etc. consist of *∼* 10^2^ of sensors that record mixtures of predominantly cortical network oscillations [1–5]. The network oscillations have distinct spatio-temporal signatures, based on the functional brain areas involved, their interconnections, and the electromagnetic mapping between the source currents and the sensors. However, the source-to-sensor mixing and the further superposition of measurement noise complicates the interpretation of the sensor-level data and its topography [6]. Given the widespread and growing availability of high-density neural recording technologies [7], there is clearly a pressing need for analysis tools that can recover underlying dynamic components from highly multivariate data.

This problem fits within a larger class of blind source separation (BSS) problems for which there are a plethora of component analysis algorithms that attempt to extract underlying source activity as linear weighted combinations of the sensor activity. The decomposition weights are designed according to some predefined criterion on the extracted time-series depending on the application. Independent Component Analysis (ICA) [8–11]) has been particularly popular within neuroscience for a number of reasons. It requires no assumption of the original sources except for statistical independence, i.e., no or minimal mutual information between the components [12], and in principle requires little to no user intervention to run. However, a drawback of the method is that it assumes that the samples of each component time trace are independent and identically distributed, which is not generally true in physical or physiological applications. This leads to another major drawback of ICA: it relies on the cumulative histograms of sensor recordings and if these histograms are Gaussian, ICA is in principle unable to separate such sources [13]. This includes, for example, physiological signals such as EEG, MEG, sEEG etc. that are at least approximately Gaussian distributed due to the central limit theorem [14], since these signals reflect the linear combination of many sources of activity. Finally, given the lack of assumptions on the structure of the underlying signals, there is no guarantee that ICA will extract relevant oscillatory components, or any other highly structured dynamic pattern for that matter.

Given the prominence and ubiquity of oscillations in neurophysiological data, applications of component analysis methods in neuroscience have taken a different approach to emphasize oscillatory dynamics, employing canonical correlation analysis [15], generalized eigenvalue decomposition [16], joint decorrelation [17], or similar methods to identify a set of spatial weights to maximize the signal-to-noise ratio in the extracted component within a narrow band around a given frequency of interest[18–21]. The extracted component then inherits the intrinsic oscillatory dynamics around that frequency without the need for a pre-designed narrow-band filter [22]. These techniques do acknowledge the inherent temporal dynamics of the oscillatory sources, but do so via nonparametric sample correlation or cross-spectrum matrix estimates that are sensitive to noise or artifacts and that require substantial amounts of data to achieve consistency.

Another problem common to all of these component decomposition methods is that they do not directly estimate the source to sensor mixing matrix. Instead, they estimate spatial filters that extract the independent components, which are not directly interpretable. The source to sensor mixing matrix can only be estimated by solving an inverse problem that requires knowledge of the noise covariance matrix [23], which adds another complication and source of potential error. Finally, the properties of the extracted component time courses are not known a priori and must be assessed after the fact, typically by using nonparametric tests as well as visual inspection.

These difficulties with component selection and interpretation occur in large part because existing blind source separation methods lack a generative model for the underlying spatio-temporal dynamics. With a probabilistic generative model, it is possible to specify a soft constraint on the dynamic properties of the underlying components, while maintaining other desirable properties such as independence between components. Component selection can be handled automatically within a statistical framework under the model, and interpretation is straightforward in principle if the components can be described by a small number of parameters. Here we propose a novel component analysis method that uses a clever state space model [24–26] to efficiently represent oscillatory dynamics in terms of latent analytic signals [27] consisting of both the real and imaginary components of the oscillation. The observed data are then represented as a superposition of these latent oscillations, each weighted by a multi-channel mixing matrix that describes the spatial signature of the oscillation. We estimate the parameters of the model and the mixing matrices using Generalized Expectation Maximization and employ empirical Bayes model selection to objectively determine the number of components. In these ways we address the major shortcomings described above for many component analysis methods. We refer to our novel method as Oscillation Component Analysis (OCA) akin to Independent Component Analysis (ICA). In what follows we describe the model formulation in detail and demonstrate the performance of the method on simulated and experimental data sets including high-density EEG during propofol anesthesia and sleep, as well as resting-state MEG from the Human Connectome Project.

## Theory

### State space oscillator model

Oscillatory time-series can be described using the following state space representation [24–26]:

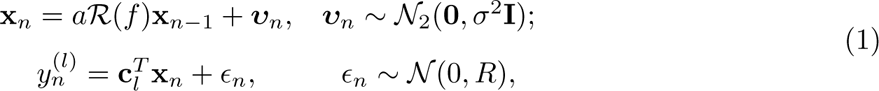

where the the oscillation state, **x***_n_* = [*x_n,_*_1_*, x_n,_*_2_]*^⊤^* is a two dimensional state vector. The stochastic difference equation summarizes oscillatory dynamics using random rotation in two-dimensional state space through a deterministic rotation matrix, *R*, explicitly parameterized by the oscillation frequency, *f* , the sampling rate, *f_s_*, and the damping factor, *a* as,

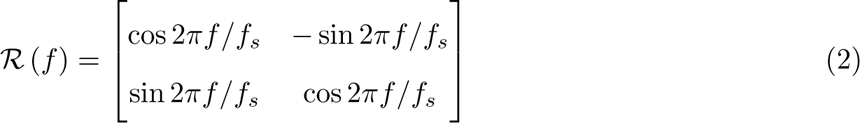

and a stochastic driving noise, assumed to be stationary and Gaussian with variance *σ*^2^. These elements of this state vector trace out two time-series that maintains an approximate *π/*2 radian phase difference, and therefore are closely related to the real and imaginary parts of an analytic signal [27] in the complex plane (see SI Text: Section Oscillation states and analytic signals ). The oscillation states are henceforth called analytic signal with minor abuse of notation. An arbitrary fixed projection (i.e., on the real line) of the state vector realizations generates the observed noisy oscillation time-series at the sensor. Multiple oscillations can be readily incorporated in this state space model by simply considering their linear combination. Recently, several investigators [28–30] have utilized this state space representation to extract underlying oscillatory time courses from single channel EEG time traces.

### Generalization to multichannel data

In order to represent multichannel neural recordings, we employ this oscillatory state space representation within a blind source separation model [31]. In the proposed generative model, a sensor-array with *L* sensors records neurophysiological signals produced by superimposition of *M* distinct oscillations supported by underlying brain networks or circuits, which we will refer to as *oscillation sources*.

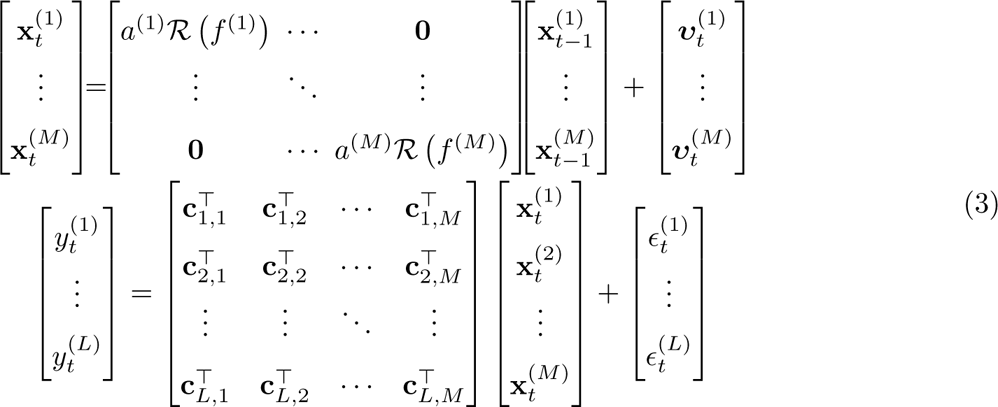

Each oscillation source is governed by the above-mentioned state space representation, resulting in a structured 2*M* dimensional state space. The structure of the underlying brain networks or circuits governs how each oscillation source is observed at the sensor array, via a *M ×* 2 spatial distribution matrix, **c***_∗,m_* = [**c**_1*,m*_, **c**_2*,m*_*, · · · ,* **c***_L,m_*]. In other words, the *l*^th^ electrode in the sensor array observes a specific projection of the underlying *m*^th^ analytic oscillation given by **c***_l,m_* = [*c_l,m,_*_1_*, c_l,m,_*_2_], which encodes the amplitude and phase of the *m*^th^ oscillation time-course at that electrode. This probabilistic generative model for multichannel recordings is motivated by a potential biophysical mechanism of electromagnetic traveling wave generation in brain parenchyma [32] (see SI Text: Section Mechanistic origin ).

### An illustration

In Fig 1(a), middle panel, we depict an oscillation state as an analytic signal, **x***_n_* (black arrows) in 2D state space that is rotating around the origin according to the given state space model of left panel: the red dashed line traces the oscillation states over time. The actual amount of rotation between every pair of consecutive time-points is a random variable centered around 2*πf/f_s_* = 0.22*π*, with the spread determined by damping parameter, *a* = 0.99 and process noise covariance. The right panel shows two noisy measurements, reflecting two different projections: one on the real axis (blue traces), another on the line making *π/*4 radian angle to the real axis (orange traces). Because of this angle between the lines of projection, these two measurements maintain approximately *π/*4 radian phase difference throughout the time course of the oscillation.

**Figure 1.**
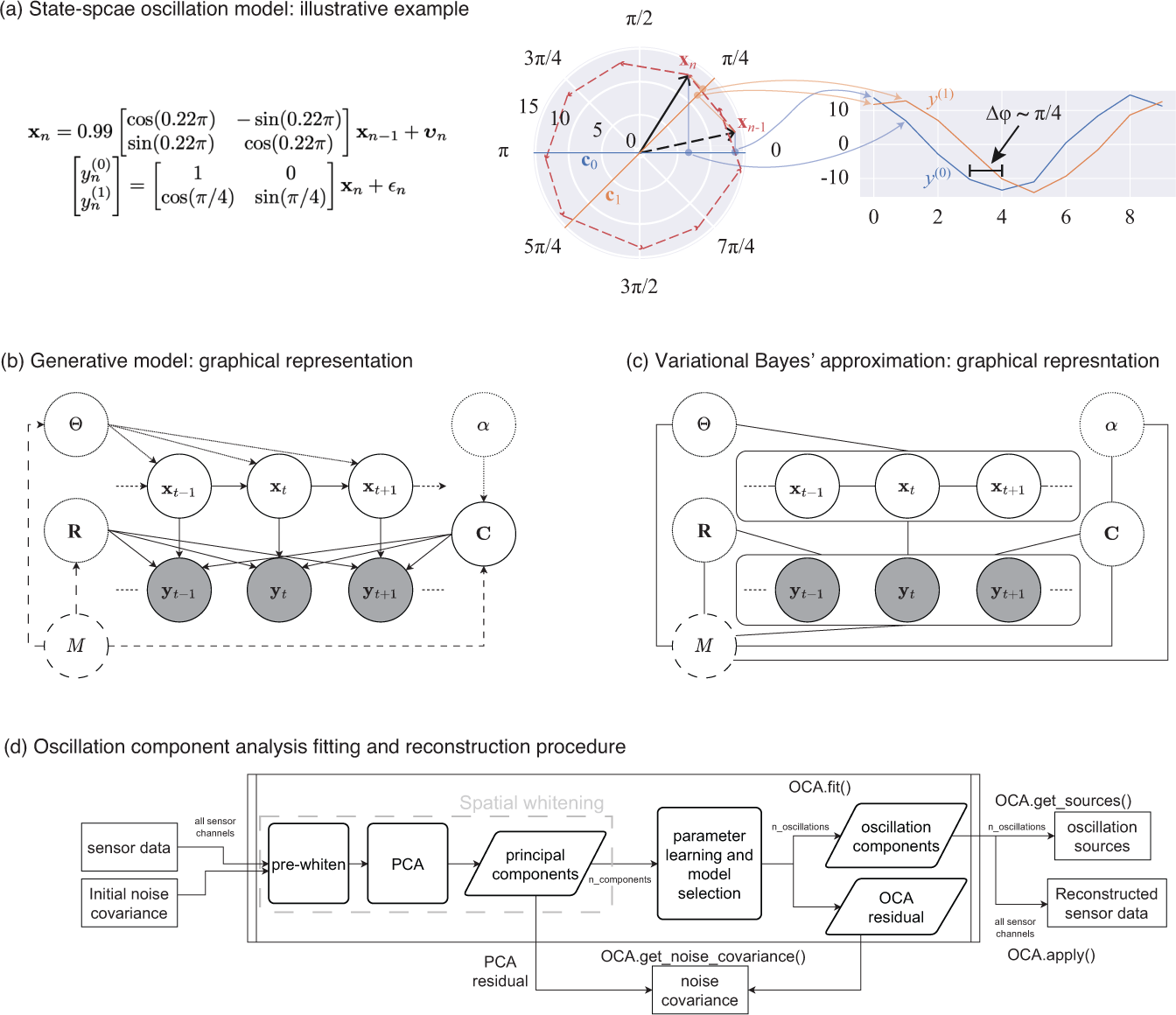
From state space oscillator model to oscillation component decomposition: (a) An illustrative example of a multichannel state space oscillation model: A single oscillation realized as an analytic signal **x***_n_* is measured as two different projections having *π/*4 radian phase difference. (b) Graphical representation of the probabilistic generative model describing the oscillations as dynamic processes, that undergo mixing at the sensors and that are observed with additive Gaussian noise. (c) Graphical representation of the Variational Bayes’ approximation that allows iterative closed form inference. (d) Oscillation component analysis fitting and reconstruction pipeline for experimentally recorded neurophysiological data. The pipeline exposes a number of methods for ease of analysis, i.e., for fitting the OCA hyperparameters fit() method, that accepts the sensor recordings and an initial sensor noise covariance matrix as input, for extracting the oscillation time-courses get sources(), for reconstructing a multichannel signal from any arbitrary subset of oscillation components, apply(), for getting a final noise covariance estimate from the residuals of OCA fitting, get noise covariance() method etc.

We note that several earlier works [33, 34] similar multivariate oscillator models, albeit from the perspective of a spatio-spectral eigendecomposition of the companion form of a multivariate autoregressive (MVAR) parameter matrix [35]. As described earlier, we employ this state space form here in the context of a blind source separation problem.

Next, we briefly describe how one can infer the hidden oscillator states from observed multivariate time-series dataset given the oscillation state space model parameters; and potentially adjust the oscillation state space model parameters to the dataset. We defer the mathematical derivations to the Materials and methods section to maintain lucidity of the presentation.

## Learning Algorithm

### Priors

We assume simple prior distributions on the measurement noise, and sensor level mixing coefficients to facilitate stable and unique recovery of the oscillation components. To obtain an *M* - oscillator analysis of *L*-channel data, we consider a Gaussian prior on the spatial mixing components, **c***_l,m_* with precision *α*, and an inverse-Wishart prior on sensor noise covariance matrix, **R** with scale, **Ψ** and degrees of freedom, *ν*:

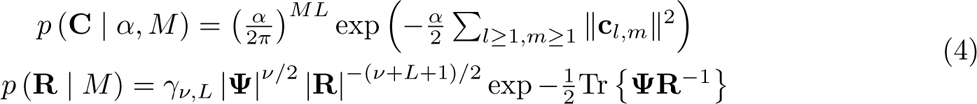

We treat the oscillation parameters, (*f* ^(*m*)^*, a*^(*m*)^, (*σ*^2^)^(*m*)^) := ***θ***^(*m*)^ and distributional parameters of the assumed priors, *α*, **Ψ***, ν* as hyperparameters. Fig 1(b) shows the probabilistic graphical model portraying oscillations as a dynamical system evolving in time, with the priors as parent nodes.

### Variational Bayes’ inference

Unfortunately, given the priors and the interplay between oscillation source time-courses and sensor-level mixing patterns, exact computation of the log-likelihood, and thus the exact posterior, is intractable. We therefore employ variational Bayes inference [36], a computationally efficient inference technique to obtain a closed-form approximation to the exact Bayes’ posterior. Originally introduced by [37], variational Bayesian inference simplifies the inference problem through a restrictive parameterization that reduces the search space for distributions, splitting up the problem into multiple partial optimization steps that are potentially easier to solve [38]. The name comes from the *negative variational free energy*, also known as evidence lower bound or ELBO [39], used to assess the quality of the aforementioned approximation and as a surrogate for the log-likelihood (see SI text: Section Negative variational free energy ).

In particular, given the number of oscillatory components *M* and the corresponding hyper-parameters ***θ*** = [***θ***^(1)^, ***θ***^(2)^*, · · · ,* ***θ***^(*M*)^] and *α,* **Ψ***, ν* we use the following Variational Bayes (VB) decoupling [38] of the posterior distribution of mixing matrix **C**, oscillation states **x***_t_* and noise covariance matrix **R**:

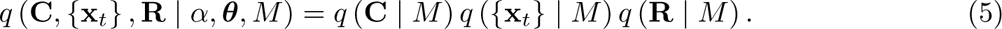

This particular choice allows the approximate posteriors of these quantities to be represented in closed-form by multivariate Gaussian, multivariate Gaussian and inverse-Wishart distributions, respectively (see SI text: Section Variational Bayes’ inference ). Fig 1(c) shows the graphical representation of posterior distribution after the VB decoupling. This essentially allows us to perform an iterative posterior inference procedure, where we cyclically update the posteriors *q* (*{***x***_t_} | M* ), *q* (**C** *| M* ), and *q* (**R** *| M* ) using the latest sufficient statistics from other two distributions (see SI text: Section Variational Bayes’ inference for more details).

### Generalized EM to update hyperparameters

Since the parameters of the state space model, and of the assumed prior distributions, are not known *a priori*, we then obtain their point estimates using an instance of the generalized Expectation Maximization (GEM) algorithm which utilizes the aforementioned approximate inference in the E-step [39, 40]. We start the learning algorithm by initializing ***θ***, 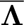, 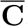, *α* to appropriate values (see SI text: Section Initialization of parameters and hyper parameters for details). We then cyclically update the posteriors *q* (*{***x***_t_} | M* ), *q* (**C** *| M* ), and *q* (**R** *| M* ) until these iterations stop changing the negative variational free energy. At the end, we update the hyperparameters ***θ***, *α* from the current inference, and proceed to the next inference with the updated hyperparameters. These update rules form an outer update loop that refines the hyperparameters, within which operates an inner update step that iteratively improves the approximate posterior distribution of the model parameters. We defer the update rules to SI text: Section Generalized EM algorithm . The algorithm terminates when the hyperparameter adjustments no longer increase the free energy of the model, i.e., we resort to parametric empirical Bayes estimation (see SI text: Section Empirical Bayes inference and model selection ).

### Selecting optimal number of oscillations

The learning algorithm assumes that the number of oscillation sources, *M* , is known a-priori, which is rarely the case for experimentally recorded data. In theory, our probabilistic treatment could assign a log-likelihood for a given dataset to every model structure, i.e., the number of oscillation sources, that can be used as a goodness-of-fit score. Unfortunately, the log-likelihood is intractable to evaluate, however the negative variational free energy can be used as a surrogate.

We formalize this idea by considering that the model structure, *M* , is drawn from a discrete uniform distribution over a finite contiguous set, (1*, M_max_*):

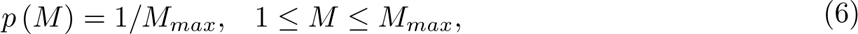

with *M_max_* being the maximal number of oscillation sources. This allows us to compute the posterior on the model structure, *q*(*M* ) within the same variational approximation framework. Using the negative variational free energy expression of the oscillator model it is easy to show that *q*(*M* ) is proportional to the exponential of the negative variational free energy of the *M* -oscillator model (see SI text: Section Model structure posterior ). Once the model posteriors are obtained, one can choose to simply select the model with maximum negative variational free energy (empirical Bayes model selection, see SI text: Section Empirical Bayes inference and model selection ), or locate the *knee* or *jump* point that exhibits a significant local change in *q*(*M* ) (for such an example see the DiffBic function[41]).

### Oscillation component analysis

A preliminary version of this algorithm has been presented previously [42]. In order to analyze experimental data, we devised the a standardized pipeline as demonstrated in Fig 1(d) closely following the MNE ICA pipeline [43]. We refer to this pipeline as OCA. The pipeline, implemented as python class OCA, accepts the sensor recordings either as continuous raw data or a set of discrete epochs, each with an initial sensor noise covariance matrix.

The hyperparameters for the measurement noise covariance prior are derived from the supplied initial sensor noise covariance matrix, and are not updated in this pipeline. The sensor data is standardized (i.e., pre-whitened) against the sensor noise covariance: this step also applies all active signal-space projectors to the data if present such as the average reference [44]. The pre-whitened data is then decomposed using PCA. The first n_components are then passed to the parameter learning iterations for hyperparameters ***θ****, α*, followed by the joint inference of the mixing matrix, oscillation time-courses, and residual noise covariance. The optimal number of oscillations is then selected as described earlier. The final noise covariance is computed from the remaining PCA components that are not supplied to oscillation fitting and the OCA residuals. Once the oscillation hyperparameters and the mixing matrix have been estimated, the oscillation time-courses can be estimated from any continuous recordings or discrete epochs. Finally, the pipeline can also reconstruct a multichannel signal from any arbitrary subset of oscillation components. We emphasize here that the main reason for using PCA here is to perform rotation of the data, so that: 1) rank-deficient data (i.e., average referenced EEG data) can be handled conveniently, by throwing out the zero-variance component. 2) the variance the multi-channel data can be uniformly distributed in the remaining PCs. For these reason, we retain components explaining 99.9% variance of the multichannel recordings, i.e., almost all components with non-zero variance. The PCA step essentially performs rotation of the data in the spatial dimension, leaving the temporal relationships untouched.

## Data Examples

Finally, we apply OCA on a simulation example and several experimentally recorded datasets. We demonstrate the versatility of OCA analysis using two EEG datasets, one under propofol-induced anesthesia, another during sleep, and one resting state MEG dataset. The simulation example shows the utility of such time-domain analysis over frequency-domain analysis in the context of multichannel recordings. In the real data applications, we showcase the implications of OCA as a mechanistically-grounded tool, suggesting various downstream analysis involving the oscillation time courses and their sensor distributions.

### Simulation Example

We first apply the proposed oscillation decomposition method on a synthetic dataset to illustrate how our time-domain approach differs from traditional frequency-domain approach. We generated a synthetic EEG dataset using a 64 channel montage, forward model and noise covariance matrix from a sample dataset distributed with the MNE software package [43]. To select the active current sources, four regions with 5 mm^2^ area were centered on chosen labels in the DKT atlas [45]: ‘transversetemporal-lh’, ‘precentral-rh’, ‘inferiorparietal-rh’ and ‘caudalmiddlefrontal-lh’. Three time-courses were separately generated at a sampling frequency of 100 Hz according to univariate AR(2) dynamics tuned to generate stable oscillations at 1.6 Hz (slow/delta), 10 Hz (alpha), 12 Hz (alpha) frequencies. The first two regions were simulated with the same slow/delta time-course such that the activity in the ‘precentral-rh’ area (orange, Fig 2A) lags that of ‘transversetemporal-lh’ area (blue, Fig 2A) by 10 ms. The last two regions, ‘inferiorparietal-rh’ and ‘caudalmiddlefrontal-lh’, were simulated with two independent alpha time-courses. We projected this source activity to the EEG sensors via a lead-field matrix and add spatially colored noise generated with the covariance structure in Fig 2D to the sensor data. Two 20 s epochs were chosen for OCA and model selection was performed within the range of (2, 3, 4, 5, 6, 8, 10) oscillations. Fig 2B presents the two most common frequency-domain visualizations of multichannel data: the top panel shows the power distribution over the scalp within different canonical EEG frequency bands, while the channel-wise power spectral density is shown in bottom panel. Fig 2C demonstrates how the decomposition obtained by OCA provides an accurate delineation of different oscillation time-courses and effective characterization of their spatial distributions in the sensor level mixing maps. Unlike the frequency-domain techniques, the residual time-series that follows OCA extraction provides the temporally unstructured part of the multichannel recording. The estimated covariance matrix from these residuals closely resembles the original sensor noise covariance matrix that was used to corrupt the recordings (see Fig 2E). The covariance matrix estimate is strikingly close to the true covariance matrix (i.e., compare with Fig 2D). OCA provides a measure, *q*(*M* ), for model structure selection, that objectively identifies the existence of 3 oscillation time-courses with different time dynamics (see Fig 2F).

**Figure 2.**
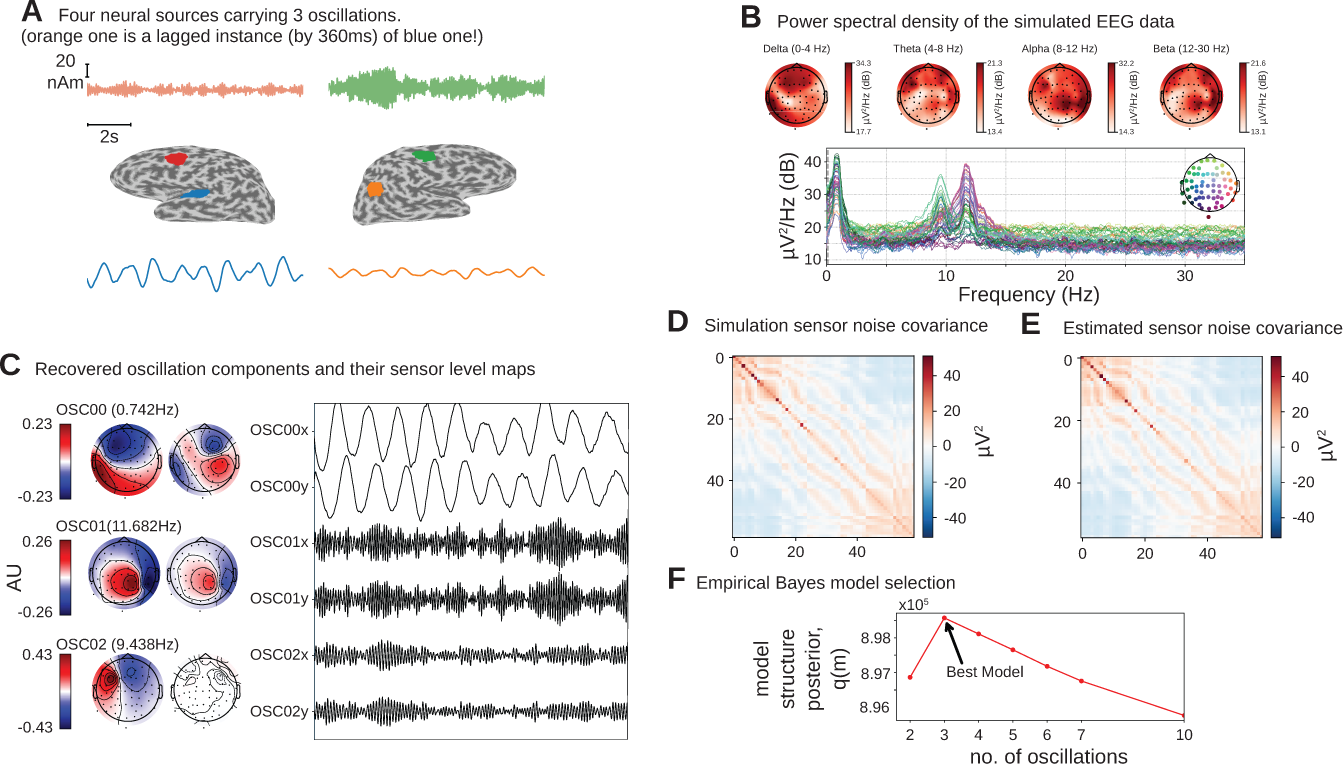
Simulation Study: **A.** four neural sources carrying 3 oscillations, where the orange time-course is a lagged instance of blue time-course, red and green time courses are independent; **B.** power spectral density of the simulated EEG recording, and power distribution over the EEG sensors in different frequency bands; **C.** recovered oscillation components and their sensor level maps; **D.** sensor noise covariance for the simulation; **E.** estimated sensor noise covariance matrix; **F.** model structure selection via model structure posterior *q*(*m*).

#### How does OCA compare to conventional approaches?

We performed additional analyses to compare OCA with Fourier based frequency domain method and ICA in a different realization of a similar synthetic EEG dataset (see Fig 3A). The frequency domain approach applied here is based on the multitaper method: given a bandwidth parameter of 2 Hz, power spectrum density is computed for individual channels. The oscillatory activity can be identified visually as the peaks in the power spectrum plot in Fig 3B top panel, and its scalp distribution can be obtained by averaging the power within given frequency bands around the peaks in each channel (as shown in Fig 3B bottom panels). However, these visualizations do very little to identify underlying oscillatory sources and pose additional questions. For example, the left topographical plot hints to two possible sources, but provides no evidence if they are separated spatially, temporally or both. Similarly, the right two topographical plots are almost identical and show no separability between these the oscillatory sources that generate two distinct peaks in spectrum. Regarding ICA, we use the ‘extended infomax’ [46] implementation provided by MNE-python 1.2 [43] to see if ICA can distinguish the three statistically independent sources in these data. The leading 4 ICA components are shown in Fig 3C. The ICA-identified components mix the underlying independent signals, which is not surprising since all components follow a Gaussian distribution. Meanwhile, OCA is able to recover three distinct oscillatory components consistent with the ground truth. OCA is able to do so because it explicitly models temporal dynamics of the components.

**Figure 3.**
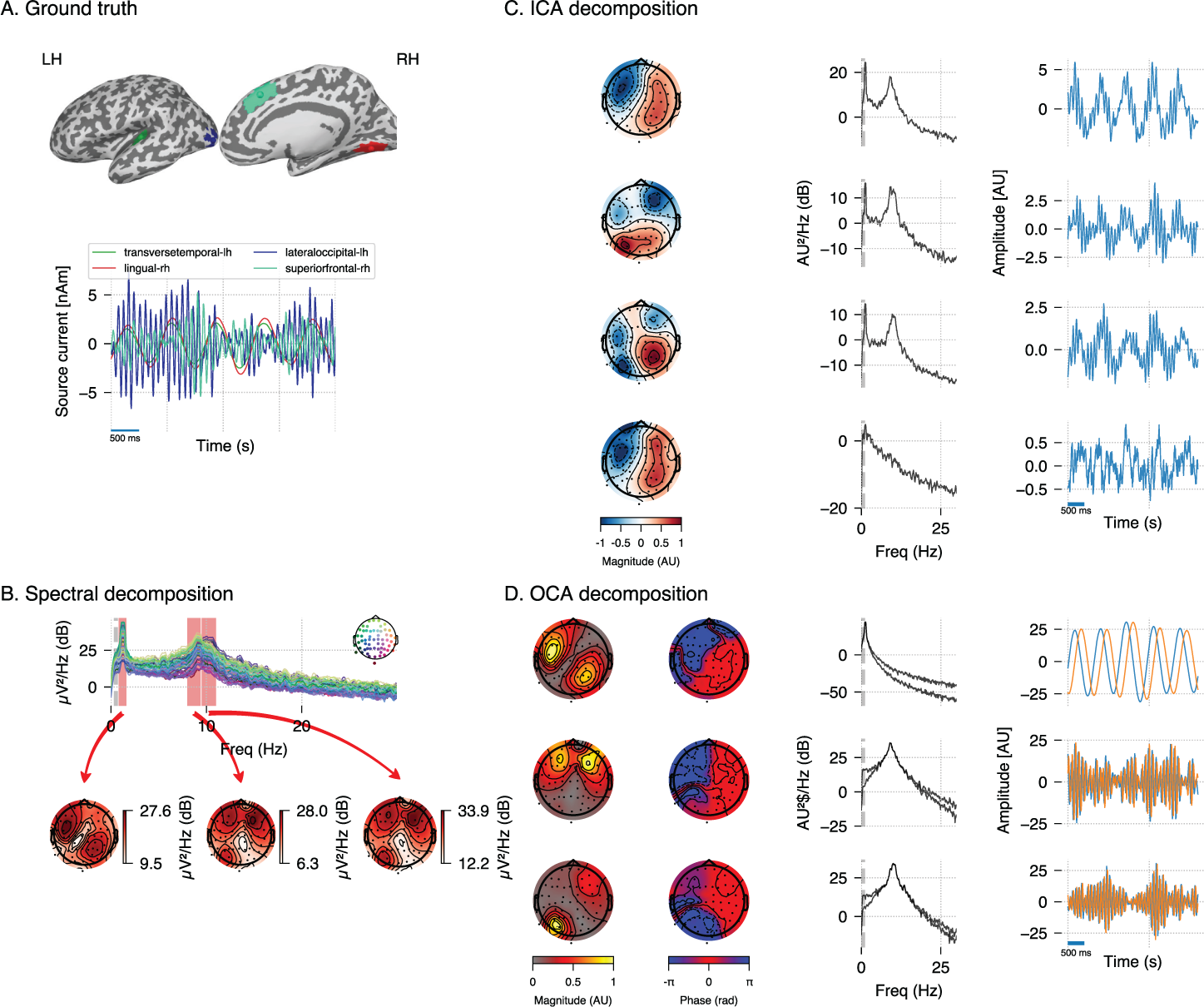
Simulation Study (extended): **A.** four neural sources carrying 3 oscillations, similar to Fig 2; **B.** power spectral density of the simulated EEG recording, and power distribution over the EEG sensors in the frequency bands (red overlay) around visually identifiable peaks; **C.** recovered ICA components (left, middle and right columns show topographic maps, power spectrum density and time-courses respectively); **D.** recovered OCA components (the topographic maps show the magnitude (left) and phase (right), while line plots show power spectrum density (left) and time-courses (right) respectively.)

We repeated similar analyses using real human EEG recording and found a similar result where ICA produced components that mixed slow and alpha band signals whereas OCA identified distinct oscillatory components (see SI text: Section Comparison of OCA to traditional approaches in experimental EEG data) .

### EEG recording during propofol-induced unconsciousness

Next, we demonstrate utility of OCA on experimental data using EEG recordings from a healthy volunteer undergoing propofol-induced unconsciousness, previously described in [47]. For induction of unconsciousness, the volunteer underwent a computer-controlled infusion of propofol to achieve monotonically increasing levels of effect-site concentration in steps of 1 µg mL*^−^*^1^. Each target effect-site concentration level was maintained for 14 min. The EEG was recorded using a 64-channel BrainVision MRI Plus system (Brain Products) with a sampling rate of 5000 Hz, bandwidth 0.016–1000 Hz. The volunteers were instructed to close their eyes throughout the study. Here we investigated EEG epochs during maintenance of effect-site concentrations of 0 µg mL*^−^*^1^ (i.e., baseline, eyes closed), 2 µg mL*^−^*^1^ and 4 µg mL*^−^*^1^ for OCA. We selected 10 clean 3.5 s epochs corresponding to those target effect site concentration (see Supplementary Materials).

We fitted OCA models with 20, 25, 30, 35, 45, 50, 55, 60 oscillation components, and selected the model with highest negative variational free-energy as the best OCA model. Following this empirical Bayes criteria we selected OCA models with 30, 30, 50 components for these three conditions, respectively. When the center frequencies were grouped within the canonical frequency bands [1], we found 17, 17, 29 slow/delta (0.1–4 Hz) oscillation components and 17, 9, 15 alpha (8–13 Hz) oscillation components for the three conditions respectively. Fig 4A–C shows power spectral densities (PSDs) of reconstructed EEG activity within each band, defined as the cumulative projection of these grouped oscillation components to the sensors. Since each of these oscillations have different sensor distributions, the increasing number of oscillation components can be associated with fragmentation of neuronal networks that support these oscillations.

**Figure 4.**
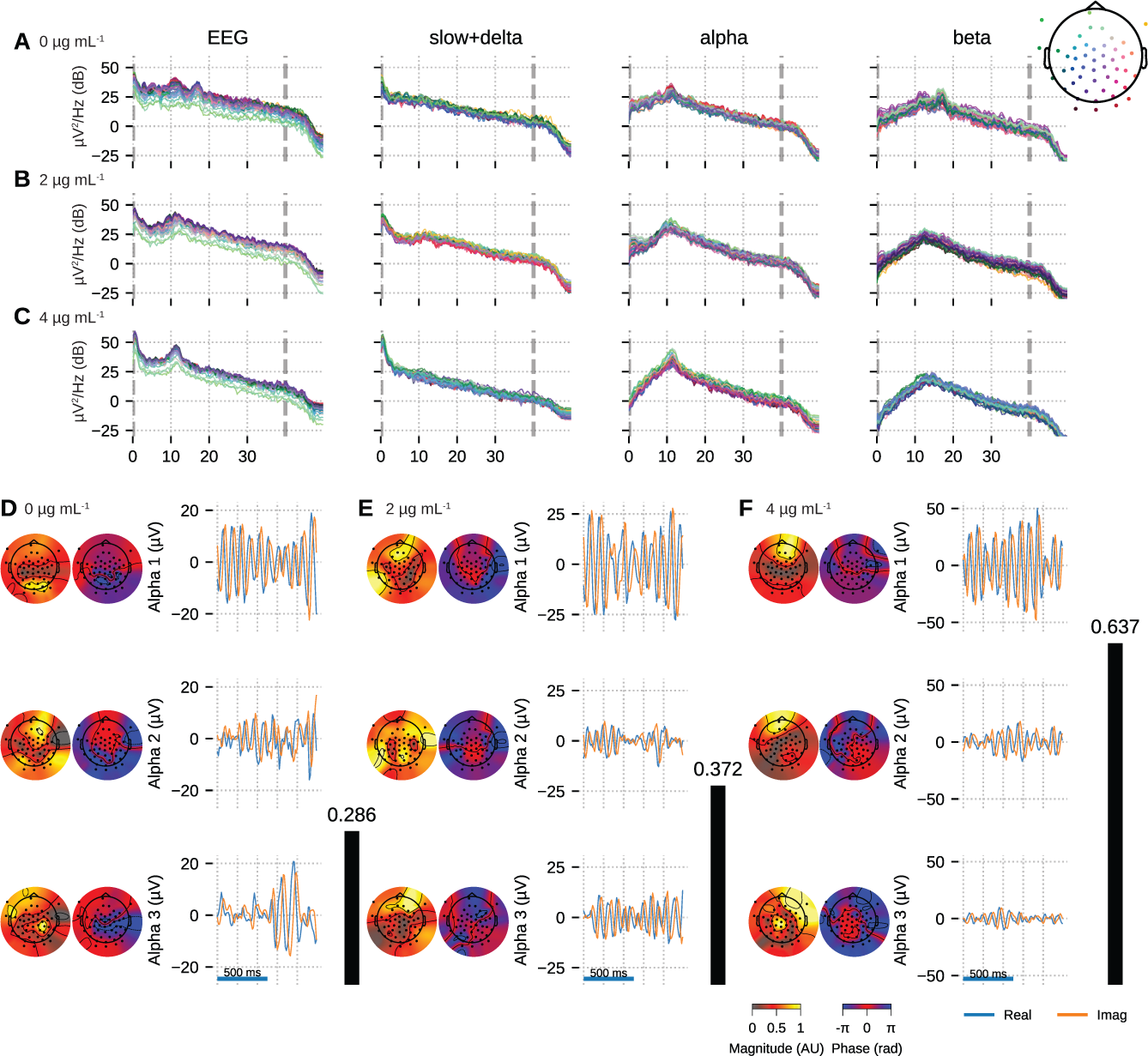
OCA of the EEG from a healthy volunteer undergoing propofol-induced unconsciousness. Conditions of target effect site concentration of **A.** & **D.** 0 (i.e. baseline), **B.** & **E.** 2 µg mL*^−^*^1^ and **C.** & **F.** 4 µg mL*^−^*^1^ are analyzed. Panels **A-C** show the PSDs of reconstructed EEG activity within each canonical band. Panels **D-F** show the three dominant (in terms of sensor wide power) alpha component: the topographic maps show the magnitude (left) and phase (right) distribution of sensor level mixing, the time courses are 1 sec representative example of the extracted oscillations from the selected epochs. The black bars on the right display the coherency measure within alpha band. OCA correctly identifies that the spatial mixing sensor maps of the alpha waves (8 Hz to 12 Hz) are oriented posteriorly at baseline, but gradually become frontally-dominant under propofol. The sensor weights are scaled to have maximum value 1. So, the units of time series traces can be considered to be in µV.

We also quantified the coherency of the reconstructed EEG activity within these commonly used frequency bands as the ratio of the power of the strongest component to the total power of the oscillation components within the bands. For the data analyzed here, slow+delta band coherency decreases gradually while the alpha band coherency increases with the increasing propofol effect-site concentration. To investigate this observation further, we visualized three dominant alpha components from each conditions in order of their strengths in Fig 4D-F. In each column, the left sub-panels consist of two topographic plots: the left one showing the distributions of the strength (scaled between 0 to 1) and the right one showing the relative phase of the estimated analytic oscillation mixing, **c***_i,j_*. The right sub-panels display examples of the extracted analytic oscillations, i.e., estimated real and imaginary components of 1 s long oscillation segments. The rightmost black bars show the coherency measure within alpha band. Clearly, during the 4 µg mL*^−^*^1^ effect site concentration, the amplitude of the leading alpha oscillation component is much bigger than the others, which aligns with our observation of the coherency measure. This finding previously reported analyses showing a rise of slow+delta oscillation power, rapid fragmentation of a slow+delta oscillation network, and emergence of global alpha network during propofol induced anesthesia [48, 49]. Further, as the propofol effect site concentration increases, the temporal dynamics increasingly resemble a pure sinusoidal signal, as evident from the PSD of the Alpha band reconstruction that becomes more concentrated around the center frequency. Lastly, the topographic plots in Fig 4D-F illustrate how alpha activity shifts from posterior electrodes to frontal electrode with increasing propofol concentration [47, 50].

### EEG recordings under different Sleep Stages

The HD-EEG data shown in Fig 5 is from a young adult subject (age 28, Female) undergoing a sleep study, recorded using ANT Neuro eego mylab system. After preprocessing (see Supplementary Materials), we visually selected 10 clean 5 s segments of data that exhibited strong alpha oscillations, each from relaxed wakeful (eyes closed) and REM sleep to apply OCA.

**Figure 5.**
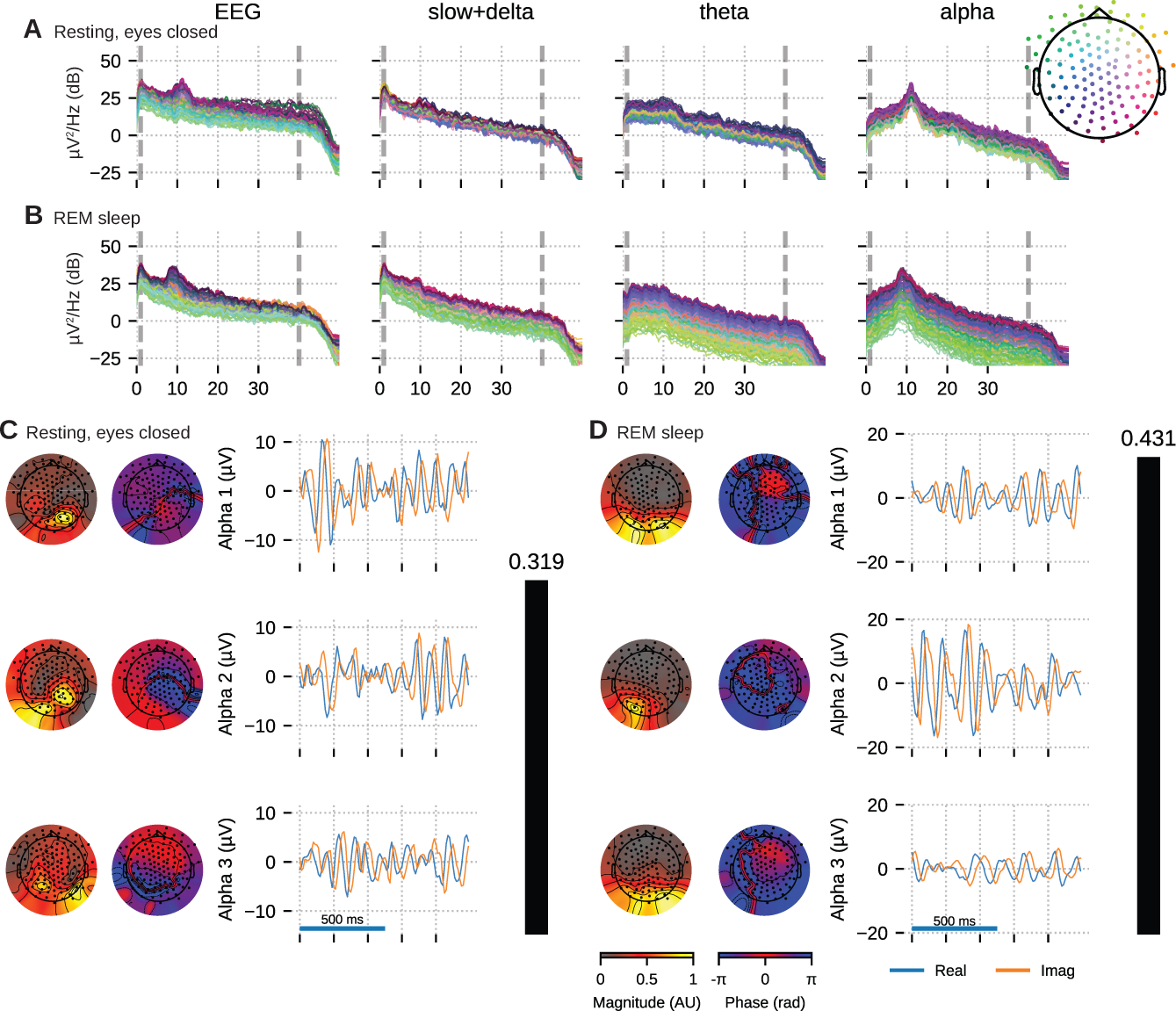
OCA of the EEG from a healthy young volunteer to compare between (**A.** & **C.**) wakeful resting state and (**B.** & **D.**) rapid eye movement sleep. Panels **A-B** show the PSDs of reconstructed EEG activity within each canonical band. Panels **C-D** show the three dominant (in terms of sensor wide power) alpha component: the topographic maps show the magnitude (left) and phase (right) distribution of sensor level mixing, the time courses are 1 sec representative example of the extracted oscillations from the selected epochs. The rightmost black bars display the coherency measure within alpha band. The contrasting topographic distribution of the alpha components (8–12 Hz), the shape of the oscillation power spectrum and alpha coherence hints at a distinct generating mechanism for alpha waves during stage–2 REM sleep compared to awake eyes closed alpha wave. The sensor weights are scaled to have a maximum value 1 so that the units of time series traces are in µV.

We fitted OCA models with 20, 25, 30, 35, 45, 50, 55, 60 oscillation components, and selected the model structure with the highest negative variational free-energy as the optimum OCA model. The empirical Bayes criterion selected 55 and 50 OCA components for relaxed wakeful (eyes closed) and REM sleep, respectively. Fig 5 follows a format similar to Fig 4: panels A-B show the power spectrum density of the recorded EEG and the reconstructed sensor data from the OCA components grouped in slow, theta and alpha bands; while panels C-D visualizes three of the extracted oscillation components within the alpha band, ordered according to their strengths. The topographical maps show the scalp distribution of these components, while the blue and red traces show 1 s–long oscillation time-series pair corresponding to each component.

Activity within the alpha band (8–13 Hz) could be summarized by a few oscillation components: for example only 7 and 3 OCA components had their center frequency within alpha band during relaxed wakeful (eyes closed) condition and REM sleep, respectively. These alpha oscillation components during relaxed wakeful (eyes closed) condition and REM sleep appeared to be distinct in terms of their temporal dynamics: relaxed wakeful alpha components are more regular (i.e. closer to sinusoidal activity) than REM alpha. But, REM alpha coherency (0.4307) is moderately higher than that of relaxed wakeful alpha (0.3195). The spatial distribution of the REM alpha component appeared to be confined primarily within the posterior channels while the awake alpha is more distributed. To quantify the degree of similarity or difference between the spatial distributions, we calculated the principal angle between the spatial mixing maps of the awake eyes-closed and REM alpha components. The *principal angle* measures the similarity between two subspaces, where an angle of 0 ° indicates that one subspace is a subset of the other, and where an angle of 90 ° indicates that at least one vector in a subspace is orthogonal to the other [51]. The spatial mixing maps for awake eyes-closed alpha and REM sleep had a principal angle of 44.31 °, suggesting that the subspaces, and thus the underlying cortical generators, were substantially different.

### Resting state MEG recording

Finally, we demonstrate the utility of OCA on a MEG dataset from the Human Connectome Project. As requested by the Human Connectome Project, the following text is copied verbatim: MEG data in Section Resting state MEG recording ‘were provided by the Human Connectome Project, WU-Minn Consortium (Principal Investigators: David Van Essen and Kamil Ugurbil; 1U54MH091657) funded by the 16 NIH Institutes and Centers that support the NIH Blueprint for Neuroscience Research; and by the McDonnell Center for Systems Neuroscience at Washington University.’ The MEG data used here is the resting-state recording from subject 104 012 MEG, session 3.

We fit OCA on 34 clean 5 s epochs, and from among the OCA with 20, 25, 30, 35, 40, 45, 50, 55 components, empirical Bayes criterion selected the OCA with 30 components. Out of these 13 were within the slow+delta band and 10 were alpha oscillations. For this example, we sought to highlight a different downstream analysis of the extracted oscillation components. We chose the three leading slow oscillation components and the three leading alpha oscillation components from the set of identified oscillation components.

For each oscillation, OCA extracts a pair of time traces, i.e., one *real* time trace and one *imaginary* time trace (see SI Text: Section Oscillation states and analytic signals )). These ‘real’ and ‘imaginary’ indices can be utilized to compute the instantaneous phase and instantaneous amplitude of individual oscillation components at every time-point, without having to rely on Hilbert transform [52].

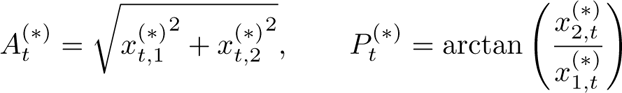

Further, we pick any one of the slow oscillation components and one of the alpha oscillation components and consider the ordered pair of slow component phase and alpha component amplitude at a single time-point as a sample drawn from the joint distribution of the ‘phase-amplitude’ of these two components. These samples then can be used to quantify cross frequency phase-amplitude coupling between these components via either nonparametric [53–55] or parametric modelling [56, 57]. Fig 6 demonstrates a possible way of investigating the coupling between the slow oscillation component phase and alpha oscillation component amplitude: the conditional mean of the alpha amplitude given slow/delta phase is higher at some specific phases between all three slow waves and OSC012, but overall flat for other alpha oscillation components. This example illustrates how the OCA algorithm can be used to identify specific oscillatory components that show phase-amplitude coupling.

**Figure 6.**
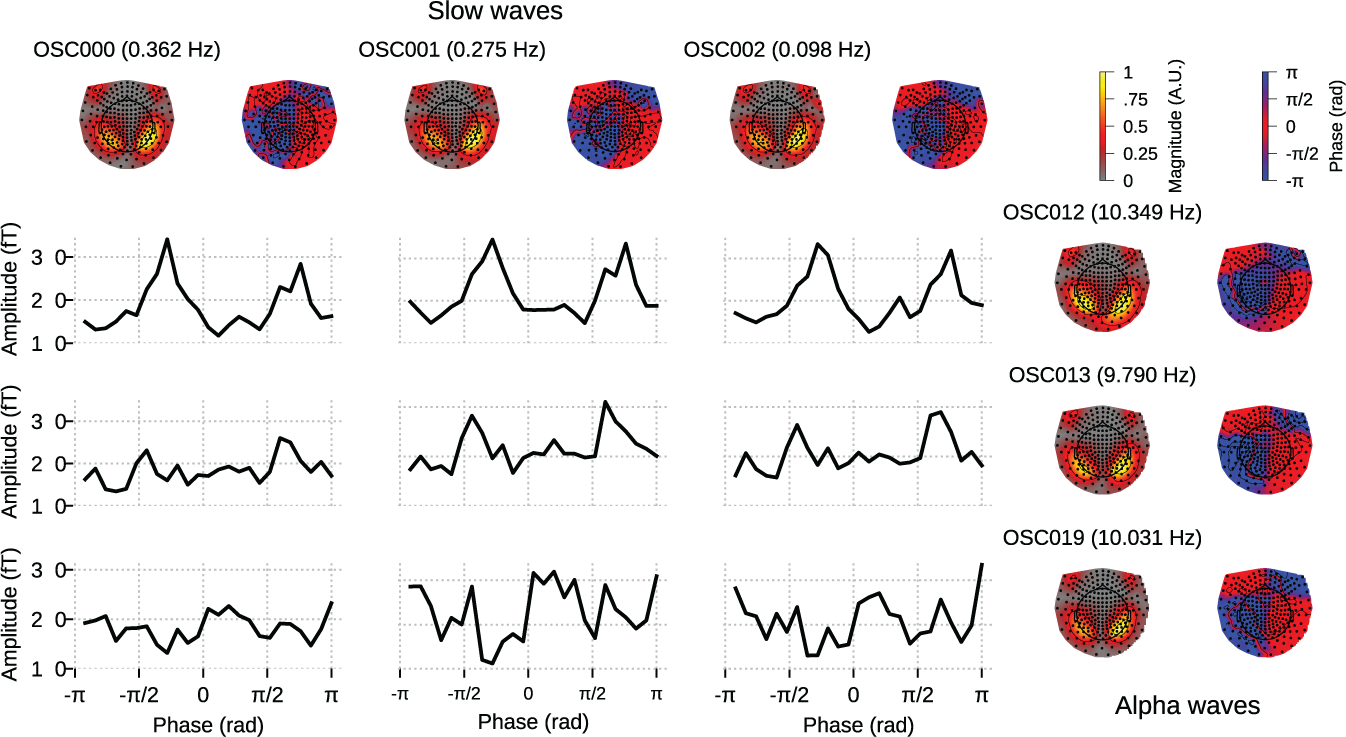
Cross–frequency phase–amplitude coupling in OCA components extracted from resting state MEG recording. The black traces show the conditional mean of a selected alpha component (8–12 Hz) amplitude given another selected slow/delta component (0–4 Hz) phase. The three slow oscillations and three alpha oscillations that explained the highest variance were selected for demonstration purposes. The topographic maps show the magnitude (left) and phase (right) distribution of sensor level mixing of the selected components.

## Discussion

OCA is a novel approach to the multichannel component decomposition problem that can identify a number of independent spatio-temporal components, but does so in the context of underlying temporal dynamics specified by a state space model. In the state space representation used here, the dynamics of an elemental oscillation are parameterized by a central frequency, *f* , a damping parameter, *a* and the second order statistics of stochastic driving noise, *σ*^2^. Meanwhile an associated spatial mixing pattern at the sensor level quantifies the contribution of the elemental oscillation to each observed channel.

The OCA learning algorithm uses an instance of the generalized expectation maximization algorithm to iteratively *match* the parameters of the oscillation state space parameters to the second order statistics of the M/EEG data. The goodness-of-fit for the data–driven matching procedure is defined within a Bayesian framework, as is the inference for the oscillation time-courses, their sensor level mixing patterns, and the measurement noise covariance matrix. This same goodness-of-fit metric is further used to determine the number of oscillations present in the multichannel data via empirical Bayes model selection. Once the number of oscillations, oscillation state space parameters, their sensor level mixing patterns, and the measurement noise covariance matrix are estimated, the elemental oscillatory activities can be extracted as a pair of time-courses, i.e., the ‘real’ and ‘imaginary’ parts of the analytical signal, from any given observation.

The oscillator state space parameters discovered in OCA are akin to the ad-hoc parameters, e.g., peak frequency, full-width-half-maxima bandwidth, and peak oscillation power etc. encountered in Fourier/wavelet based frequency-domain nonparametric time series analysis [58]. However, OCA circumvents a major drawback of frequency-domain methods when searching for neural oscillations from multichannel recordings. Generally, the source-to-sensor mixing complicates the interpretation of the topographies generated by signal processing tools that treat each channel individually [6]. In a nonparametric frequency-domain approach, the only way to discover this spatially-correlated structure across sensors is to perform eigenvalue analysis on the cross-spectral density on a frequency by frequency basis, also known as global coherence analysis [50, 59, 60]. If the eigenvalue distribution exhibits sufficient skewness, the associated frequency is deemed coherent and the leading eigenvectors are identified as the ‘principal’ sensor networks at that frequency; in other words a proxy for a strong oscillatory component. As the number of sensors increases this analysis becomes increasingly intractable: in a sensor array with *∼* 10^2^ sensors the cross–spectral density matrix has *∼* 10^4^ entries. Alternatively, the state space representation used here in OCA provides a convenient and compact way to perform multichannel time-domain modeling and analysis [61]. The OCA modelling approach decouples the oscillatory dynamics and the spatial mixing pattern effectively by estimating one set of parameters for each discovered oscillation and the associated spatial mixing matrix, thereby avoiding the need for cross–spectral density matrices and frequency parameters for individual channels altogether [62]. Another major advantage of OCA over global coherence analysis is that OCA automatically identifies the dominant *coherent* oscillations across the channels based on its probabilistic generative model and Bayesian learning approach.

The iterative parameter estimation procedure of OCA is clearly more computationally burdensome compared to conventional Frequency-domain nonparametric methods. However, once the OCA parameters are estimated, OCA can decompose any given multichannel recording segments into the oscillation time-series pairs. In that sense, OCA parameter learning can be viewed as iterative estimation of a pair of optimal spatio–temporal filters [63] for each oscillation, parameterized by the estimated oscillation state space parameters and spatial maps. These optimal spatio–temporal filters are then applied to the multichannel data to extract the oscillation time courses. As a result, the extracted narrowband activity is endogenous to the data, rather than being imposed by an arbitrary narrowband filter [22].

This behavior of OCA is similar to oscillatory component extraction methods based on data-driven spatial filtering, where the extracted component inherits the intrinsic oscillatory dynamics around the specified frequency [18, 19, 19, 20]. In fact, OCA employs the same philosophy as spatio-temporal source separation [21], where the extracted narrow-band signal and its time-shifted versions are again projected to a temporal coordinate space to further enhance the power within the given frequency band. However, spatio-temporal source separation establishes the temporal constraint via nonparametric sample correlation estimates that are sensitive to noise or artifacts and that require substantial amounts of data for estimation. Another important distinction is that spatio-temporal source separation determines the sensor weights first, and then obtains the temporal filter kernel from the projection of the multichannel data. In contrary, OCA updates the sensor weights and the parameters for the temporal filter iteratively within the Bayesian formulation of the state space framework, thus jointly estimating the spatial and temporal components simultaneously from the data.

The class of ICA–based methods, on the other hand, assumes that the component time courses share no mutual information, i.e., are statistically independent. Without any explicit assumption on the temporal dynamics of the generative process, the properties of the identified components may be difficult to interpret and may also be unreliable. In fact, the identified ICA components typically require subjective visual inspection by experts for their possible interpretation. A number of investigators have recognized these limitations with ICA, in particular the assumption of temporal independence, and have proposed generalizations of the ICA framework to incorporate auto-regressive modeling of source dynamics [31, 64]. Despite demonstrating improved source separation performance in naturalistic signals like music [64], adoption of these methods for the analysis of neurophysiological data has been slow. This may be due in part to the lack of interpretability of the higher order auto-regressive model structures. Alternatively, Brookes et. al. use a combination of Hilbert envelope computation within pre-defined frequency bands and temporal down-sampling to identify meaningful temporally independent time signals, but stop short of trying to disambiguate different oscillatory sources [13]. Thus, they only assess the connectivity pattern within pre-defined bands, i.e., how different areas of the brain are harmonized through modulation of the oscillations or vice-versa inside those pre-defined bands. The spatial maps recovered from anatomically projected resting state MEG data by this method resemble spatial patterns of fMRI resting state networks. OCA is close to the first variant of ICA, but describes the identified component sources using a two-dimensional state space vector that efficiently represents oscillatory dynamics [28] in a manner that is easy to interpret. Matsuda and Komaki [33] described a similar state space model for multivariate time series, but did not apply the model to neurophysiological data, perhaps due to its high dimensionality. The OCA algorithm differs by way of its learning and model selection algorithms, which allow it to select the number of components to extract in a statistically principled and data driven manner. Overall, OCA delivers the best of the previously mentioned eigen-analysis-based source separation methods and ICAs: a decomposition method that can identify independent components, but where each component represents an underlying oscillatory dynamical system. The generative link between each component and the underlying dynamical system makes interpretation of the components straightforward. Here we focus specifically on analyzing neurophysiological data that are exemplified by highly structured oscillatory dynamics. But the methods we describe apply to more general, arbitrary dynamics that can be approximated by linear state space models, and can be equally useful so long as the state space model is interpretable.

There is another important distinction between OCA and the aforementioned blind source separation techniques. These methods directly estimate the sensor weights for a weighted linear combination of sensor recordings that produce the individual temporal components [23]. In other words, these blind source separation methods provide a backward decoding algorithm. The linear weights are not directly interpretable since they do not specify how the individual temporal components map to the observed data. That uniquely interpretable information is provided by a forward mapping which must be estimated in a separate step. When the number of linear components matches with the number of sensors, this forward mapping can be calculated via simple matrix inversion. However, when there is a reduced set of components the forward mapping can only be obtained by solving an inverse problem that requires knowledge of the source and sensor sample covariance matrix. In contrast, the OCA model involves only the (forward) spatial distribution matrix and our algorithm directly estimates it. In fact, the backward extraction model of OCA involves non-zero weights for time-shifted signals and are never explicitly computed. In that sense, OCA avoids an extraneous transformation step that could inadvertently introduce errors to the spatial mixing pattern estimates.

Another popular time-domain approach for oscillatory signal discovery is the multichannel extension of empirical mode decomposition. Empirical mode decomposition models the time series data as linear combination of intrinsic oscillations called intrinsic mode functions (IMF). Identification of IMFs depends critically on finding the local mean of the local extrema, which is not well-defined in the context of multichannel recordings. Rehman and Mandic [65] chose to take real-valued projections of the *n*-channel data, with the projections taken along direction vectors uniformly sampled on a unit spherical surface in an *n* dimensional coordinate space. The local extrema of each of the projected signals are extrapolated to obtain a set of multivariate signal envelopes. Clearly, this random sampling in a high-dimensional space is computationally demanding, and the fully nonparametric formulation requires a substantial amount of data, making the procedure sensitive to noise. On the contrary, OCA employs a parametric model that represents oscillatory dynamics in a manner that is statistically efficient and resilient to noise.

## Conclusion

In summary, starting from a simple probabilistic generative model of neural oscillations, OCA provides a novel data-driven approach for analyzing multichannel synchronization within underlying oscillatory modes that are easier to interpret than conventional frequency-wise, cross-channel coherence. The overall approach adds significantly to existing methodologies for spatio-temporal decomposition by adding a formal representation of dynamics to the underlying generative model. The application of OCA on simulated and real M/EEG data demonstrates its capabilities as a principled dimensionality reduction tool that simultaneously provides a parametric description of the underlying oscillations and their activation pattern over the sensor array.

## Materials and methods

### M/EEG pre-processing

All prepossessing were performed using MNE-python 1.2[43] and Eelbrain 0.37[66], with default setting of the respective functions.

#### EEG recording during propofol-induced unconsciousness

The EEG was bandpass filtered between 0.1 Hz to 40 Hz, followed by downsampling to 100 Hz and average referencing prior to performing OCA. The 0.1 Hz highpass filtering was done to filter out slow drifts in the EEG recordings, which can adversely effect the OCA fitting procedure. All selected epochs had peak-to-peak signal amplitude less than 1000mV. The prior on the noise-covariance was estimated from the same EEG recording, after high-pass filtering above 30 Hz.

#### Sleep EEG recording

The EEG data was first down sampled to 100 Hz after band pass filtering within 1 Hz to 40 Hz. The flat and noisy channels are first identified upon visual inspection. We computed neighborhood correlation for the rest of the channels and marked channels with median correlation, *ρ <* 0.4 as bad channels. These channels are dropped for the subsequent analysis.

#### MEG recording from Human Connectome Project

We divided the entire resting state MEG recording from subject 104 012 MEG, session 3 into 5 s epochs, and selected 34 epochs with peak-to-peak amplitude less than 4000 fT for OCA. Prior to fitting OCA, we used Signal-space projection [44] to remove heartbeat and eye movement artifacts from MEG signals. The repaired MEG recordings are then downsampled to 100 Hz after applying appropriate anti-aliasing filter.

## Supporting Information

**Supporting Text: Mechanistic origin** Traveling wave solution of Maxwell’s equation to the state space model.

**Supporting Text: Oscillation states and analytic signals** Relation between oscillation states and analytic signals

**Supporting Text: Model parameter estimation and model selection** Detailed derivation of the OCA algorithm.

**Supporting Text: Comparison of OCA to traditional approaches in experimental EEG data**

## Acknowledgments

This work was supported by National Institutes of Health (Grant no. R01AG054081-01A1) and Tiny Blue Dot Foundation.

## References

1. G. Buzsaki, Rhythms of the Brain. Oxford University Press, 2006.

2. G. Buzsáki and A. Draguhn, “Neuronal Oscillations in Cortical Networks,” Science, vol. 304, pp. 1926–1929, June 2004.

3. F. Lopes da Silva, “EEG and MEG: Relevance to Neuroscience,” Neuron, vol. 80, pp. 1112– 1128, Dec. 2013.

4. X.-J. Wang, “Neurophysiological and Computational Principles of Cortical Rhythms in Cognition,” Physiological Reviews, vol. 90, pp. 1195–1268, July 2010.

5. R. F. Helfrich and R. T. Knight, “Oscillatory Dynamics of Prefrontal Cognitive Control,” Trends in Cognitive Sciences, vol. 20, pp. 916–930, Dec. 2016.

6. N. Schaworonkow and V. V. Nikulin, “Is sensor space analysis good enough? Spatial patterns as a tool for assessing spatial mixing of EEG/MEG rhythms,” NeuroImage, p. 119093, Mar. 2022.

7. I. H. Stevenson and K. P. Kording, “How advances in neural recording affect data analysis,” Nature Neuroscience, vol. 14, pp. 139–142, Feb. 2011.

8. A. Hyvärinen and E. Oja, “Independent component analysis: Algorithms and applications,” Neural Networks, vol. 13, pp. 411–430, June 2000.

9. T.-P. Jung, S. Makeig, C. Humphries, T.-W. Lee, M. J. McKeown, V. Iragui, and T. J. Sejnowski, “Removing electroencephalographic artifacts by blind source separation,” Psychophysiology, vol. 37, pp. 163–178, Mar. 2000.

10. J.-F. Cardoso, “High-Order Contrasts for Independent Component Analysis,” Neural Computation, vol. 11, pp. 157–192, Jan. 1999.

11. A. Hyvärinen, “Independent component analysis: Recent advances,” Philosophical Transactions of the Royal Society A: Mathematical, Physical and Engineering Sciences, vol. 371, p. 20110534, Feb. 2013.

12. A. J. Bell and T. J. Sejnowski, “An Information-Maximization Approach to Blind Separation and Blind Deconvolution,” Neural Computation, vol. 7, pp. 1129–1159, Nov. 1995.

13. M. J. Brookes, M. Woolrich, H. Luckhoo, D. Price, J. R. Hale, M. C. Stephenson, G. R. Barnes, S. M. Smith, and P. G. Morris, “Investigating the electrophysiological basis of resting state networks using magnetoencephalography,” Proceedings of the National Academy of Sciences, vol. 108, pp. 16783–16788, Oct. 2011.

14. R. J. Muirhead, Aspects of Multivariate Statistical Theory. Wiley Series in Probability and Mathematical Statistics, New York: Wiley, 1982.

15. N. Robinson, K. P. Thomas, and A. P. Vinod, “Canonical correlation analysis of EEG for classification of motor imagery,” in 2017 IEEE International Conference on Systems, Man, and Cybernetics (SMC), (Banff, AB), pp. 2317–2321, IEEE, Oct. 2017.

16. L. Parra and P. Sajda, “Blind Source Separation via Generalized Eigenvalue Decomposition,” Journal of Machine Learning Research, vol. 4, pp. 1261–1269, 2003.

17. A. de Cheveigné and L. C. Parra, “Joint decorrelation, a versatile tool for multichannel data analysis,” NeuroImage, vol. 98, pp. 487–505, Sept. 2014.

18. V. V. Nikulin, G. Nolte, and G. Curio, “A novel method for reliable and fast extraction of neuronal EEG/MEG oscillations on the basis of spatio-spectral decomposition,” NeuroImage, vol. 55, pp. 1528–1535, Apr. 2011.

19. A. de Cheveigné and D. Arzounian, “Scanning for oscillations,” Journal of Neural Engineering, vol. 12, p. 066020, Dec. 2015.

20. M. X. Cohen, “Multivariate cross-frequency coupling via generalized eigendecomposition,” eLife, vol. 6, p. e21792, Jan. 2017.

21. M. X. Cohen, “Using spatiotemporal source separation to identify prominent features in multichannel data without sinusoidal filters,” European Journal of Neuroscience, vol. 48, pp. 2454–2465, Oct. 2018.

22. N. Yeung, R. Bogacz, C. B. Holroyd, S. Nieuwenhuis, and J. D. Cohen, “Theta phase resetting and the error-related negativity,” Psychophysiology, vol. 44, pp. 39–49, 2007.

23. S. Haufe, F. Meinecke, K. Görgen, S. Dähne, J.-D. Haynes, B. Blankertz, and F. Bießmann, “On the interpretation of weight vectors of linear models in multivariate neuroimaging,” NeuroImage, vol. 87, pp. 96–110, Feb. 2014.

24. N. Wiener, Nonlinear Problems in Random Theory. Aug. 1966.

25. A. C. Harvey, Forecasting, Structural Time Series Models, and the Kalman Filter. Cambridge; New York: Cambridge University Press, 1989.

26. A. C. Harvey and T. M. Trimbur, “General Model-Based Filters for Extracting Cycles and Trends in Economic Time Series,” Review of Economics and Statistics, vol. 85, pp. 244–255, May 2003.

27. R. N. Bracewell, The Fourier Transform and Its Applications. McGraw-Hill Series in Electrical and Computer Engineering, Boston: McGraw Hill, 3rd ed ed., 2000.

28. T. Matsuda and F. Komaki, “Time series decomposition into oscillation components and phase estimation,” Neural Computation, vol. 29, no. 2, pp. 332–367, 2017.

29. A. M. Beck, E. P. Stephen, and P. L. Purdon, “State Space Oscillator Models for Neural Data Analysis,” in 2018 40th Annual International Conference of the IEEE Engineering in Medicine and Biology Society (EMBC), (Honolulu, HI), pp. 4740–4743, IEEE, July 2018.

30. A. M. Beck, M. He, R. Gutierrez, and P. L. Purdon, “An iterative search algorithm to identify oscillatory dynamics in neurophysiological time series,” preprint, Neuroscience, Nov. 2022.

31. L. C. Parra, “Temporal models in blind source separation,” in Adaptive Processing of Sequences and Data Structures (J. G. Carbonell, J. Siekmann, G. Goos, J. Hartmanis, J. Van Leeuwen, C. L. Giles, and M. Gori, eds.), vol. 1387, pp. 229–247, Berlin, Heidelberg: Springer Berlin Heidelberg, 1998.

32. V. L. Galinsky and L. R. Frank, “Universal theory of brain waves: From linear loops to nonlinear synchronized spiking and collective brain rhythms,” Physical Review Research, vol. 2, p. 023061, Apr. 2020.

33. T. Matsuda and F. Komaki, “Multivariate Time Series Decomposition into Oscillation Components,” Neural Computation, vol. 29, pp. 2055–2075, Aug. 2017.

34. A. J. Quinn, G. G. Green, and M. Hymers, “Delineating between-subject heterogeneity in alpha networks with Spatio-Spectral Eigenmodes,” NeuroImage, vol. 240, p. 118330, Oct. 2021.

35. A. Neumaier and T. Schneider, “Estimation of parameters and eigenmodes of multivariate autoregressive models,” ACM Transactions on Mathematical Software, vol. 27, pp. 27–57, Mar. 2001.

36. A. Quinn and V. Šmídl, The Variational Bayes Method in Signal Processing. Signals and Communication Technology Ser, New York Boulder: Springer NetLibrary, Inc., 2006.

37. G. E. Hinton and D. van Camp, “Keeping the neural networks simple by minimizing the description length of the weights,” in Proceedings of the Sixth Annual Conference on Computational Learning Theory - COLT ’93, (Santa Cruz, California, United States), pp. 5–13, ACM Press, 1993.

38. H. Attias, “Inferring Parameters and Structure of Latent Variable Models by Variational Bayes,” in Proceedings of the Fifteenth Conference on Uncertainity in Artificial Intelligence, pp. 21–30, July 1999.

39. R. M. Neal and G. E. Hinton, “A View of the EM Algorithm that Justifies Incremental, Sparse, and other Variants,” in Learning in Graphical Models (M. I. Jordan, ed.), pp. 355–368, Dordrecht: Springer Netherlands, 1998.

40. A. P. Dempster, N. M. Laird, and D. B. Rubin, “Maximum Likelihood from Incomplete Data Via the *EM* Algorithm,” Journal of the Royal Statistical Society: Series B (Methodological), vol. 39, pp. 1–22, Sept. 1977.

41. Q. Zhao, M. Xu, and P. Fränti, “Knee Point Detection on Bayesian Information Criterion,” in 2008 20th IEEE International Conference on Tools with Artificial Intelligence, (Dayton, OH, USA), pp. 431–438, IEEE, Nov. 2008.

42. P. Das and P. L. Purdon, “Extracting common oscillatory time-courses from multichannel recordings: Oscillation Component Analysis,” in 2022 56th Asilomar Conference on Signals, Systems, and Computers, (Pacific Grove, CA, USA), pp. 602–606, IEEE, Oct. 2022.

43. A. Gramfort, M. Luessi, E. Larson, D. A. Engemann, D. Strohmeier, C. Brodbeck, L. Parkkonen, and M. S. Hämäläinen, “MNE software for processing MEG and EEG data,” NeuroImage, vol. 86, pp. 446–460, Feb. 2014.

44. M. A. Uusitalo and R. J. Ilmoniemi, “Signal-space projection method for separating MEG or EEG into components,” Medical & Biological Engineering & Computing, vol. 35, pp. 135–140, Mar. 1997.

45. R. S. Desikan, F. Ségonne, B. Fischl, B. T. Quinn, B. C. Dickerson, D. Blacker, R. L. Buckner, A. M. Dale, R. P. Maguire, B. T. Hyman, M. S. Albert, and R. J. Killiany, “An automated labeling system for subdividing the human cerebral cortex on MRI scans into gyral based regions of interest,” NeuroImage, vol. 31, pp. 968–980, July 2006.

46. T.-W. Lee, M. Girolami, and T. J. Sejnowski, “Independent Component Analysis Using an Extended Infomax Algorithm for Mixed Subgaussian and Supergaussian Sources,” Neural Computation, vol. 11, pp. 417–441, Feb. 1999.

47. P. L. Purdon, E. T. Pierce, E. A. Mukamel, M. J. Prerau, J. L. Walsh, K. F. Wong, A. F. Salazar-Gomez, P. G. Harrell, A. L. Sampson, A. Cimenser, S. Ching, N. J. Kopell, C. Tavares-Stoeckel, K. Habeeb, R. Merhar, and E. N. Brown, “Electroencephalogram signatures of loss and recovery of consciousness from propofol,” Proc Natl Acad Sci U S A, vol. 110, pp. E1142–51, Mar. 2013.

48. L. D. Lewis, V. S. Weiner, E. A. Mukamel, J. A. Donoghue, E. N. Eskandar, J. R. Madsen, W. S. Anderson, L. R. Hochberg, S. S. Cash, E. N. Brown, and P. L. Purdon, “Rapid fragmentation of neuronal networks at the onset of propofol-induced unconsciousness,” Proceedings of the National Academy of Sciences, vol. 109, Dec. 2012.

49. M. Murphy, M.-A. Bruno, B. A. Riedner, P. Boveroux, Q. Noirhomme, E. C. Landsness, J.-F. Brichant, C. Phillips, M. Massimini, S. Laureys, G. Tononi, and M. Boly, “Propofol Anesthesia and Sleep: A High-Density EEG Study,” Sleep, vol. 34, pp. 283–291, Mar. 2011.

50. A. Cimenser, P. L. Purdon, E. T. Pierce, J. L. Walsh, A. F. Salazar-Gomez, P. G. Harrell, C. Tavares-Stoeckel, K. Habeeb, and E. N. Brown, “Tracking brain states under general anesthesia by using global coherence analysis,” Proceedings of the National Academy of Sciences, vol. 108, pp. 8832–8837, May 2011.

51. A. Bjorck and G. H. Golub, “Numerical Methods for Computing Angles Between Linear Subspaces,” Mathematics of Computation, vol. 27, p. 579, July 1973.

52. A. Wodeyar, M. Schatza, A. S. Widge, U. T. Eden, and M. A. Kramer, “A state space modeling approach to real-time phase estimation,” eLife, vol. 10, p. e68803, Sept. 2021.

53. M. X. Cohen, “Assessing transient cross-frequency coupling in EEG data,” Journal of Neuroscience Methods, vol. 168, pp. 494–499, Mar. 2008.

54. N. Cheng, Q. Li, S. Wang, R. Wang, and T. Zhang, “Permutation Mutual Information: A Novel Approach for Measuring Neuronal Phase-Amplitude Coupling,” Brain Topography, vol. 31, pp. 186–201, Mar. 2018.

55. R. Martínez-Cancino, J. Heng, A. Delorme, K. Kreutz-Delgado, R. C. Sotero, and S. Makeig, “Measuring transient phase-amplitude coupling using local mutual information,” NeuroImage, vol. 185, pp. 361–378, Jan. 2019.

56. H. Soulat, E. P. Stephen, A. M. Beck, and P. L. Purdon, “State space methods for phase amplitude coupling analysis,” Scientific Reports, vol. 12, p. 15940, Sept. 2022.

57. A. Perley and T. P. Coleman, “A Mutual Information Measure of Phase-Amplitude Coupling using High Dimensional Sparse Models,” in 2022 44th Annual International Conference of the IEEE Engineering in Medicine & Biology Society (EMBC), (Glasgow, Scotland, United Kingdom), pp. 21–24, IEEE, July 2022.

58. T. Donoghue, M. Haller, E. J. Peterson, P. Varma, P. Sebastian, R. Gao, T. Noto, A. H. Lara, J. D. Wallis, R. T. Knight, A. Shestyuk, and B. Voytek, “Parameterizing neural power spectra into periodic and aperiodic components,” Nature Neuroscience, vol. 23, pp. 1655–1665, Dec. 2020.

59. V. S. Weiner, D. W. Zhou, P. Kahali, E. P. Stephen, R. A. Peterfreund, L. S. Aglio, M. D. Szabo, E. N. Eskandar, A. F. Salazar-Gomez, A. L. Sampson, S. S. Cash, E. N. Brown, and P. L. Purdon, “Propofol disrupts alpha dynamics in distinct thalamocortical networks underlying sensory and cognitive function during loss of consciousness,” preprint, Neuroscience, Apr. 2022.

60. P. Mitra and H. Bokil, Observed Brain Dynamics. Oxford ; New York: Oxford University Press, 2008.

61. N. S. Nise, Control Systems Engineering. Hoboken, NJ: Wiley, 6th ed ed., 2011.

62. H. Gunasekaran, L. Azizi, V. Van Wassenhove, and S. K. Herbst, “Characterizing endogenous delta oscillations in human MEG,” Scientific Reports, vol. 13, p. 11031, July 2023.

63. B. D. O. Anderson and J. B. Moore, Optimal Filtering. Dover Books on Engineering, Mineola, NY: Dover Publ, dover ed., unabridged republ ed., 2005.

64. B. Pearlmutter and L. Parra, “Maximum likelihood blind source separation: A context-sensitive generalization of ICA,” in Advances in Neural Information Processing Systems (M. Mozer, M. Jordan, and T. Petsche, eds.), vol. 9, MIT Press, 1996.

65. N. Rehman and D. P. Mandic, “Multivariate empirical mode decomposition,” Proceedings of the Royal Society A: Mathematical, Physical and Engineering Sciences, vol. 466, pp. 1291–1302, May 2010.

66. C. Brodbeck, T. L. Brooks, P. Das, S. Reddigari, and J. P. Kulasingham, “Christianbrodbeck/Eelbrain: 0.37.” Zenodo, Apr. 2022.

