## Supporting Info for "A dynamic generative model can extract interpretable oscillatory components from multichannel neurophysiological recordings"

### Supporting information

#### Supporting text

##### Mechanistic origin

Here we detail how the multichannel multi-oscillator state space model can be understood in terms of a biophysical model of brain wave generation.

**Charge-continuity and electromagnetic brain waves** Maxwell's equation provides the following charge-continuity equation that governs the spontaneous spatial-temporal evolution of the potential, $\Phi$ in brain tissues:

$$\frac{\partial}{\partial t}\boldsymbol{\nabla}^2\Phi = -\boldsymbol{\nabla}\cdot\boldsymbol{\Sigma}\cdot\Phi. \tag{S1.1}$$

This differential equation is satisfied by traveling waves of the form, $\Phi \sim \exp\left[-j\left(\mathbf{k}\cdot\mathbf{r}+\Omega t\right)\right]$, where $\mathbf{r}$ and $t$ denote the spatial coordinate and time respectively with the wave number, $\mathbf{k}$ and the complex frequency, $\Omega$ satisfying following dispersion relation: $D(\Omega,\mathbf{k}) = j\Omega|k|^2 - \Sigma_{ij}k_ik_j - j\partial_i\Sigma_{ij}k_j$, in tensor notation. Separating the real and imaginary parts of the complex frequency, $\Omega = \omega + j\gamma$, we arrive at following expressions for decaying and oscillatory frequencies:

$$\gamma = -\frac{\Sigma_{ij}k_ik_j}{|\mathbf{k}|^2}, \qquad \omega = \frac{\partial_i\Sigma_{ij}k_j}{|\mathbf{k}|^2}. \tag{S1.2}$$

Clearly, wave solutions $\boldsymbol{\Phi}$ satisfying $|\gamma/\omega| \ll 1$, i.e., the oscillatory frequency is much larger than the decaying frequency, can give rise to electromagnetic oscillations in brain parenchyma, governed by the brain composition and architecture. Detailed treatment of such a model, considering the anisotropy and inhomogeneity of the brain tissue, can be found in [1].

**Traveling wave solution to state space model** Here we consider one such oscillatory electric potential being recorded at $L$ sensors placed at coordinates $\mathbf{r}_1, \mathbf{r}_2, \cdots, \mathbf{r}_L$ (w.r.t a reference electrode placed at infinity). The potentials sampled (sampling frequency, $f_s = 1/\Delta t$) at those electrodes at time index $n$ are given as:

$$\Phi(l,n) = \Re\left\{A_l \exp\left[-j\left(\mathbf{k}\cdot\mathbf{r}_l + \Omega\, n\Delta t\right)\right]\right\} \tag{S1.3}$$

with $A_l$ representing the amplitude observed at $l$-th electrode. The separability of the spatial part $\phi^{spatial}(l) = A_l \exp\left[-j\left(\mathbf{k} \cdot \mathbf{r}_l\right)\right]$ and the temporal part $\Phi^{temporal}(n) = \exp\left[-j\left(\Omega\ n\Delta t\right)\right]$ of the general solution is crucial for the following development, since only the temporal part, $\Phi^{temporal}(n+1)$ evolves in time,

$$\Phi^{temporal}(n+1) = \exp\left[j\Omega\Delta t\right]\Phi^{temporal}(n). \tag{S1.4}$$

, while the spatial part $\phi^{spatial}(l)$ remains constant. Adapting the following real-valued matrix-vector notation that consists of the real and imaginary parts of the temporal and spatial components, $\Phi^{temporal}(n) = x_{n,1} + jx_{n,2}$ and $\Phi^{spatial}(l) = m_{l,1} + jm_{l,2}$, we arrive at:

$$\begin{bmatrix} x_{n+1,1} \\ x_{n+1,2} \end{bmatrix} = \exp\left[-\gamma\Delta t\right] \begin{bmatrix} \cos(\omega\Delta t) & -\sin(\omega\Delta t) \\ \sin(\omega\Delta t) & \cos(\omega\Delta t) \end{bmatrix} \begin{bmatrix} x_{n,1} \\ x_{n,2} \end{bmatrix}, \tag{S1.5}$$

$$y^{(l)}_{n+1} = \begin{bmatrix} A_l m_{l,1} & A_l m_{l,2} \end{bmatrix} \begin{bmatrix} x_{n+1,1} \\ x_{n+1,2} \end{bmatrix}, \tag{S1.6}$$

where $y^{(l)}_{n+1}$ is the recorded potential at electrode $l$ at time index $(n+1)$. Essentially, the complex valued oscillatory function, $\Phi^{temporal}(n)$ is an analytic signal whose real and imaginary parts constitute the ordered pair, $\mathbf{x}_n = [x_{n,1}, x_{n,2}]$, which we call the *oscillation state*. Equation (S1.5) describes a circular motion of the oscillation state $\mathbf{x}_n$ in a 2D state space with frequency $\omega$ at every time step through the damped rotation matrix $\exp\left[-\gamma\Delta t\right]\mathcal{R}(\omega\Delta t)$, while a projection, dictated by the electrode location (Eq S1.6), is recorded at the EEG electrodes as an oscillation (see Fig 1(a)). This deterministic model forms the basis for the probabilistic linear Gaussian state space model in 1 which introduces process and observation noise, $\boldsymbol{v}_n$ and $\epsilon_n$ respectively, with redefined spatial components as $c_{l,*} = A_l m_{l,*}$, and the decay term $\exp\left[-\gamma\Delta t\right]$ as a damping parameter, $a$. We also represented the frequency of oscillation, $f$ as f $= \omega/2\pi$ by replacing $\omega\Delta t = 2\pi f/f_s$.

##### Oscillation states and analytic signals

Here, we consider real valued observations, $y_n \in \mathbb{R}$ admitting a state space representation in complex plane, i.e., generated from complex valued states, $z_n \in \mathbb{C}$:

$$\begin{aligned} z_n &= \rho x_{n-1} + q_n \\ y_n &= \tfrac{1}{2}(z_n + \bar{z}_n) + r_n \end{aligned} \tag{S2.1}$$

where $\bar{z}_n$ denotes complex conjugate of $\bar{z}_n$, $q_n$ and $r_n$ are complex-valued and real-valued random variables. In stead-state, i.e., $t \to \infty$, the states, $z_n$ admit Fourier representation given by:

$$Z(e^{-j\omega}) = \frac{Q(e^{-j\omega})}{1 - \rho e^{-j\omega}} \tag{S2.2}$$

It is clear that if $q_n$ is an analytic signal, i.e., $Q(e^{-j\omega})$ has no negative frequency component, the same will be true for $Z(e^{-j\omega})$.

Further, since $q_n$ is an analytic signal, there exists a real valued signal $s_n$, such that $q_n = s_n + j(h * s)_n$ where discrete time Fourier transform of $h_n$ is given by:

$$H(e^{-j\omega}) = \begin{cases} -j & \text{for } 0 < \omega \leq \pi \\ 0 & \text{for } \omega = 0 \\ j & \text{for } -\pi < \omega < 0 \end{cases} . \tag{S2.3}$$

Considering $s_n$ to be a white noise with variance $\sigma^2$, it is easy to verify that $h_n * s_n$ is also white noise with variance $\sigma^2$, and uncorrelated to $s_n$. This makes $q_n$ to be a 'analytic' white noise, i.e., power spectrum density of $q_n$ is flat for $0 > \omega > \pi$, and zero in $-\pi > \omega > 0$. We can then derive the following state space representation, with real valued but two dimensional states as in Eq 1, but with additional constraint that $\upsilon_{n,1}$ and $\upsilon_{n,2}$ are related by: $\upsilon_{n,2} = (h * \upsilon_1)_n$. But, since such constraint is not easy to enforce during the inference or model learning from experimental data, so we relaxed the constraint on the state-noise covariance to be the one stated in the Eq 1.

**Extraction of instantaneous amplitude and phase.** However, we can still compute the instantaneous amplitude and phase of the signal from this state space representation from the following observation. The oscillation states assume their successive values in a way, that traces limit cycle like maps around origin (see Fig 1(a)). This allows us to define the instantaneous phase as the angle the oscillation state makes from a fixed reference direction, at any given time point [2, section A.1 & 2]). For simplicity, we consider the positive direction on the real line as the reference. The amplitude is simply given by the distance from the origin.

$$A_t^{(*)} = \sqrt{{x_{t,1}^{(*)}}^2 + {x_{t,2}^{(*)}}^2} \qquad P_t^{(*)} = \arctan\left(\frac{x_{2,t}^{(*)}}{x_{1,t}^{(*)}}\right) \tag{S2.4}$$

In this sense, the oscillator state space representation provides a generalization of phasor concept, similar to analytic signals (i.e., by allowing for time-varying amplitude, phase and frequency, in contrary to invariant amplitude, phase and frequency of phasor) but with relatively relaxed constraint on the relation between real and imaginary part.

##### Model parameter estimation and model selection

**Negative variational free energy** For a $L$-channel M/EEG recording $\mathbf{y}_t$, $t = 1, 2, \cdots, T$, $M$-oscillator probabilistic state space oscillator model admits to the following distribution:

$$p\left(\{\mathbf{x}_t, \mathbf{y}_t\} \mid \mathbf{C}, \mathbf{R}, \mathbf{F}, \mathbf{Q}, M\right) = \prod_{t=1}^{T} \frac{1}{\sqrt{|(2\pi)\mathbf{R}|}} \exp -\frac{1}{2} \left\|\mathbf{y}_t - \mathbf{C}\mathbf{x}_t\right\|^2_{\mathbf{R}^{-1}} \\ \frac{1}{\sqrt{|(2\pi)\mathbf{Q}|}} \exp -\frac{1}{2} \left\|\mathbf{x}_t - \mathbf{F}\mathbf{x}_{t-1}\right\|^2_{\mathbf{Q}^{-1}} \tag{S3.1}$$

Since $\mathbf{F}$ and $\mathbf{Q}$ are parameterized using hyperparameter $\boldsymbol{\theta}$, we will re-parameterize left hand side of Eq S3.1 as $p\left(\{\mathbf{x}_t, \mathbf{y}_t\} \mid \mathbf{C}, \mathbf{R}, \boldsymbol{\theta}, M\right)$. To simplify our exposition, we define $\mathbf{X} = [\mathbf{x}_1, \mathbf{x}_2, \cdots, \mathbf{x}_N]$ and use $\mathbf{X}$ and $\{\mathbf{x}_t\}$ interchangeably (similarly for $\mathbf{Y}$). We also extensively use $vec(\circ)$ notation and associated terminology introduced in [3, page 17].

We note that computation of ensemble likelihood of the presented model requires marginalizing over $\mathbf{X}, \mathbf{C}, \mathbf{R}$ and $M$ given the priors in Eq 4 and Eq 6:

$$\begin{aligned} p\left(\{\mathbf{y}_t\}\right) &= \sum_{M=1}^{M_{max}} \int p\left(\mathbf{X}, \mathbf{Y}, \mathbf{C}, \mathbf{R} \mid \alpha, \boldsymbol{\theta}, M\right) p(M) d\left\{\mathbf{X}, \mathbf{C}, \mathbf{R}\right\} \\ &= \sum_{M=1}^{M_{max}} \int p\left(\mathbf{X}, \mathbf{Y} \mid \mathbf{C}, \mathbf{R}, \alpha, \boldsymbol{\theta}, M\right) p\left(\mathbf{C} \mid \alpha, M\right) p\left(\mathbf{R}\right) p\left(M\right) d\left\{\mathbf{X}, \mathbf{C}, \mathbf{R}\right\}. \end{aligned}$$

However, this involves an intractable integration that cannot be performed analytically. We avoid this intractability by using a lower bound to the ensemble likelihood as a surrogate for the same. Specifically, we invoke Neal-Hinton representation theorem [4] to obtain a lower bound on the ensemble log-likelihood as:

$$\begin{aligned} \mathcal{L} &= \log p\left(\{\mathbf{y}_t\}\right) \\ &= \log \sum_{M=1}^{M_{max}} \int p\left(\mathbf{X}, \mathbf{Y}, \mathbf{C}, \mathbf{R} \mid \alpha, \boldsymbol{\theta}, M\right) p(M) d\left\{\mathbf{X}, \mathbf{C}, \mathbf{R}\right\} \\ &= \log \sum_{M=1}^{M_{max}} \int \frac{p\left(\mathbf{X}, \mathbf{Y}, \mathbf{C}, \mathbf{R} \mid \alpha, \boldsymbol{\theta}, M\right) p(M)}{q\left(\mathbf{X}, \mathbf{C}, \mathbf{R} \mid M\right) q(M)} q\left(\mathbf{X}, \mathbf{C}, \mathbf{R} \mid M\right) q(M) d\left\{\mathbf{X}, \mathbf{C}, \mathbf{R}\right\} \end{aligned}$$

$$
\geq \sum_{M=1}^{M_{max}} q(M) \int \log \frac{p\left(\mathbf{X},\mathbf{Y},\mathbf{C},\mathbf{R} \mid \alpha,\boldsymbol{\theta},M\right)p(M)}{q\left(\mathbf{X},\mathbf{C},\mathbf{R},\mid M\right)q(M)} q\left(\mathbf{X},\mathbf{C},\mathbf{R} \mid M\right) d\left\{\mathbf{X},\mathbf{C},\mathbf{R}\right\}
$$

$$
= \sum_{M=1}^{M_{max}} q\left(M\right)\left[\log \frac{p\left(M\right)}{q\left(M\right)} + \int \log \frac{p\left(\mathbf{X},\mathbf{Y},\mathbf{C},\mathbf{R},\alpha,\boldsymbol{\theta} \mid M\right)}{q\left(\mathbf{X},\mathbf{C},\mathbf{R} \mid M\right)} q\left(\mathbf{X},\mathbf{C},\mathbf{R} \mid M\right) d\left\{\mathbf{X},\mathbf{C},\mathbf{R}\right\}\right]
$$

$$
= \left\langle \log \frac{p\left(M\right)}{q\left(M\right)} + \left\langle \log \frac{p\left(\mathbf{X},\mathbf{Y},\mathbf{C},\mathbf{R},\alpha,\boldsymbol{\theta} \mid M\right)}{q\left(\mathbf{X},\mathbf{C},\mathbf{R} \mid M\right)} \right\rangle_{q(\mathbf{X},\mathbf{C},\mathbf{R}|M)} \right\rangle_{q(M)} := \mathcal{F}, \tag{S3.2}
$$

which holds for any arbitrary conditional distribution $q$ and can be computed analytically if $q$ is chosen carefully. This lower bound is known as *negative variational free energy*[5], and attains the actual ensemble log-likelihood only when $q$ is the exact Bayes' posterior distribution [6]. In the last line, we use $\langle \circ \rangle_{q(\mathbf{X},\mathbf{C},\mathbf{R}|M)}$ to denote the average of the expression within $\langle\rangle$ with respect to the model posterior $q\left(\mathbf{X},\mathbf{C},\mathbf{R} \mid M\right)$.

Since negative variational free energy as a lower bound on the ensemble log-likelihood, maximization of negative variational free energy will tend to maximize of the ensemble log-likelihood. Even though global maximization of the ensemble log-likelihood is not guaranteed, the ensemble log-likelihood at the point of maximum negative variational free energy is guaranteed to at least greater than the negative variational free energy. On the other hand, the negative variational free energy can be shown to be proportional to the negative Kullback-Liebler divergence between the exact posterior and approximate posterior [5]. So, the maximization shall result in closer approximation exact posterior. In the next section, we work with the intuition that maximization of negative variational free energy leads to better approximation to the exact posterior and increment of ensemble log-likelihood.

**Variational Bayes' inference** However, the fact that the ensemble log-likelihood cannot be found analytically, makes computation of the exact Bayes' posterior distribution intractable. We thus employ an efficient variational Bayes inference procedure that approximates the exact Bayes' posterior with a distribution of form:

$$
q\left(\left\{\mathbf{x}_t\right\},\mathbf{C},\mathbf{R} \mid M\right) = q\left(\left\{\mathbf{x}_t\right\} \mid M\right) q\left(\mathbf{C} \mid M\right) q\left(\mathbf{R} \mid M\right), \tag{S3.3}
$$

i.e. that factorizes over $\left\{\mathbf{x}_t\right\},\mathbf{C},\mathbf{R}$, conditional on the model structure, $M$. This particular choice of functional constraint leads to tractable conditional distributions when the trailing term of Eq

S3.2 is subjected to maximization w.r.t. functional form:

$$q(\mathbf{X} \mid M) = \frac{1}{\sqrt{|(2\pi)\mathbf{\Sigma}_x|}} \exp -\frac{1}{2} \|vec(\mathbf{X}) - \boldsymbol{\mu}_x\|^2_{\mathbf{\Sigma}_x{}^{-1}} \quad \text{(S3.4)}$$

$$q(\mathbf{C} \mid M) = \frac{1}{\sqrt{|(2\pi)\mathbf{\Sigma}_c|}} \exp -\frac{1}{2} \|vec(\mathbf{C}) - \boldsymbol{\mu}_c\|^2_{\mathbf{\Sigma}_c{}^{-1}} \quad \text{(S3.5)}$$

$$q(\mathbf{R} \mid M) = \gamma_{\rho,L} |\mathbf{\Omega}|^{\rho/2} |\mathbf{R}|^{-(\rho+L+1)/2} \exp -\frac{1}{2}\mathrm{Tr}\left\{\mathbf{\Omega}\mathbf{R}^{-1}\right\} \quad \text{(S3.6)}$$

Here we omit the trivial derivation of the functional form for brevity of the presentation, curious readers are encouraged to expand the trailing term of Eq S3.2, and follow [6, Section 2.4] as an exercise to obtain the optimal functional forms.

Once the above-mentioned optimal functional forms of the approximate posterior distribution have been identified, next task is to find the optimal quantities that parametrize those functional forms. We show that the parameters of any of these distribution can be expressed in terms of the M/EEG data and moments of other two distributions. Generally speaking, the expressions can be found by collecting relevant terms in the expansion of trailing term of Eq S3.2, followed by simple algebraic manipulation. For example, the parameters of $q(\mathbf{R} \mid M)$ are given by:

$$\mathbf{\Omega} = \mathbf{\Psi} + \sum_{t=1}^{T} \langle \mathbf{y}_t - \mathbf{C}\mathbf{x}_t \rangle_{q(\mathbf{X},\mathbf{C}|M)} \text{ and } \rho = \nu + L. \quad \text{(S3.7)}$$

Expression for the parameters of the Gaussian distribution is a bit involved if one attempts collecting terms, and try completing squares. Instead we use the following facts about Gaussian log-posteriors: 1) since the mode and mean of a multivariate Gaussian distribution coincides, the mean of Gaussian distributions can be found by maximizing the approximate log-posterior, and 2) the negative Hessian of the log-posterior is the inverse of covariance matrix [7]. It is worth mentioning here that the log-posterior maximization can be carried out in closed form by solving a high-dimensional system of linear equations (first order optimality criterion, [8, pp. 140]), which involves Hessian matrix inversion.

In case of the latent oscillator states, q($\mathbf{X} \mid M$), the expressions of inverse covariance matrix and the mean vector is as follows:

$$\Sigma_x^{-1} = \begin{bmatrix} \mathbf{D} & \mathbf{S} & \mathbf{0} & \mathbf{0} & \cdots & \mathbf{0} & \mathbf{0} \\ \mathbf{S}^\top & \mathbf{D} & \mathbf{S} & \mathbf{0} & \cdots & \mathbf{0} & \mathbf{0} \\ \mathbf{0} & \mathbf{S}^\top & \mathbf{D} & \mathbf{S} & \cdots & \mathbf{0} & \mathbf{0} \\ \vdots & \vdots & \vdots & \vdots & \vdots & \vdots & \vdots \\ \mathbf{0} & \mathbf{0} & \mathbf{0} & \mathbf{0} & \cdots & \mathbf{S}^\top & \mathbf{D} \end{bmatrix} \qquad \begin{aligned} &\text{where} \\ \mathbf{D} &= \left\langle \mathbf{C}^\top \mathbf{R}^{-1} \mathbf{C} \right\rangle_{q(\mathbf{C},\mathbf{R}|M)} \\ &\quad + \widehat{\mathbf{F}}^\top \widehat{\mathbf{Q}}^{-1} \widehat{\mathbf{F}} + \widehat{\mathbf{Q}}^{-1} \\ \mathbf{S} &= -\widehat{\mathbf{F}}^\top \widehat{\mathbf{Q}}^{-1} \end{aligned} \tag{S3.8}$$

$$\boldsymbol{\mu}_x = \Sigma_x \times vec\left(\mathbf{B}\mathbf{y}_t\right) \qquad \mathbf{B} = \left\langle \mathbf{C}^\top \mathbf{R}^{-1} \right\rangle_{q(\mathbf{C},\mathbf{R}|M)}$$

We exploit the block tri-diagonal structure of the hessian matrix to realize a fast and stable inverse [9, 10]. This facilitates fast computation of the mean and covariance matrix of the approximate log-posterior.

Before tackling the case of the mixing matrix, $\mathbf{C}$, we note that each latent oscillator state is a pair of two coordinates and these two oscillator coordinates correspond to two consecutive columns of the mixing matrix, i.e., contribution of any given latent oscillator state on the observation can be viewed as the inner product, i.e., $\mathbf{y}_t^{(l)} = \mathbf{c}_{l,m}\mathbf{x}_t^{(m)}$, so that the column pairs ($\mathbf{C}_{:,2l-1:2l}$) and latent oscillator states ($\mathbf{X}_{2l-1:2l,:}$) are only unique up to a scaling and a rotation. We resolve this ambiguity by simply fixing the first row of the mixing matrix column pair to $\begin{bmatrix} 1 & 0 \end{bmatrix}$ for all the oscillators. With this modification, the parameters of $q(\mathbf{C} \mid M)$ are given as:

$$\begin{aligned} \Sigma_c^{-1} &= \boldsymbol{\Lambda}_{2:L,2:L} \otimes T\mathbf{P}_{xx} + \alpha \mathbf{I}_{2(L-1)M\times 2(L-1)M} \\ \boldsymbol{\mu}_c &= \Sigma_c \left( T\text{vec}\left( \left( \boldsymbol{\Lambda}_{2:N}\mathbf{P}_{yx} \right)^\top \right) - \boldsymbol{\lambda}_1 \otimes \left( \mathbf{P}_{xx}\mathbf{m}_1^\top \right) \right) \end{aligned} \tag{S3.9}$$

where

$$\boldsymbol{\Lambda} = \left\langle \mathbf{R}^{-1} \right\rangle_{q(\mathbf{R}|M)}, \ \mathbf{P}_{xx} = \frac{1}{T}\sum_{t=1}^{T} \left\langle \mathbf{x}_t\mathbf{x}_t^\top \right\rangle_{q(\mathbf{X}|M)}, \ \mathbf{P}_{yx} = \frac{1}{T}\sum_{t=1}^{T} \left\langle \mathbf{y}_t\mathbf{x}_t^\top \right\rangle_{q(\mathbf{X}|M)}.$$

Using the properties of Kronecker products[3, pp 73-76], the inversion of the negative Hessian matrix and solution of the system of the linear equations can be carried out in a fast and efficient manner.

Evidently, this interconnectedness of the distributional parameters of the marginals of $\{\mathbf{x}_t\}, \mathbf{C}, \mathbf{R}$, given model structure $M$, is a testament to the inter-dependence of these three variables. Since joint posterior distribution of is assumed to factorize over the marginals, i.e., $\{\mathbf{x}_t\}, \mathbf{C}, \mathbf{R}$, given model structure $M$ and observations $\mathbf{Y}$ are assumed to be independent, inter-dependence has been captured by the interrelated distributional parameters. In fact, the interrelated parameters pass information among them using the sufficient statistic (SS) of these distributions, that are required to compute the averages in the update equations. The VB approach for posterior inference therefore leads to an iterative scheme for each model structure, $M$ when the hyperparameters $(\boldsymbol{\theta}, \alpha)$ are specified: we cyclically update the marginal posterior parameters of $\{\mathbf{x}_t\}, \mathbf{C}, \mathbf{R}$ according to Eqs. S3.8, S3.9, S3.7 respectively. Each time a distribution (i.e. its parameters) is updated, the SS associated with the distribution is also recomputed to pass to the next update. The iterations are repeated until a stopping criteria is met: stopping criteria could be a fixed number of iterations or when the objective, i.e., variational free energy stabilizes. The marginal posterior parameters obtained at the last iteration are the optimal distributional parameters providing the variational Bayes' inference.

**Generalized EM algorithm** The variational Bayes' inference of form presented in S3.3 still requires the hyperparameters $\boldsymbol{\theta}$ and $\alpha$ to be specified. The hyperparameters that maximizes the negative variational free energy are the second best choice for the hyperparameters, right after the ones that maximize the ensemble likelihood. Since the direct maximization w.r.t. the hyperparameters is impractical, we use an generalized version of Expectation maximization (EM) algorithm that uses the variational posterior inference instead of the exact posterior to compute the expectation of the complete log-likelihood expression [7, 11, 12] to learn these hyperparameters from the M/EEG recording. One requires the following averages computed w.r.t. the approximate posterior $q(\mathbf{X} \mid M)$,

$$\begin{aligned}
\mathbf{U}^{(m)} &= \textstyle\sum_{t=1}^{T} \left\langle \mathbf{x}_{t-1}^{(m)} \mathbf{x}_{t-1}^{(m)\top} \right\rangle_{q(\mathbf{X}|M)}, \\
\mathbf{V}^{(m)} &= \textstyle\sum_{t=1}^{T} \left\langle \mathbf{x}_{t.t-1}^{(m)} \mathbf{x}_{t,t-1}^{(m)\top} \right\rangle_{q(\mathbf{X}|M)}, \\
\mathbf{W}^{(m)} &= \textstyle\sum_{t=1}^{T} \left\langle \mathbf{x}_{t}^{(m)} \mathbf{x}_{t}^{(m)\top} \right\rangle_{q(\mathbf{X}|M)},
\end{aligned}$$

to update the hyperparameters $\boldsymbol{\theta}^{(m)} = \left(f^{(m)},\ a^{(m)},\ (\sigma^2)^{(m)}\right)$ as follows:

$$f^{(m)} = \frac{1}{2\pi} \arctan \left\{ \frac{\mathrm{rt}\left\{\mathbf{V}^{(m)}\right\}}{\mathrm{tr}\left\{\mathbf{V}^{(m)}\right\}} \right\}, \tag{S3.10}$$

$$a^{(m)} = \frac{\sqrt{\mathrm{tr}\left\{\mathbf{V}^{(m)}\right\}^2 + \mathrm{rt}\left\{\mathbf{V}^{(m)}\right\}^2}}{\mathrm{tr}\left\{\mathbf{U}^{(m)}\right\}}, \tag{S3.11}$$

$$\sigma^{2(m)} = \frac{1}{2T} \left( \mathrm{tr}\left\{\mathbf{W}^{(m)}\right\} - a^{(m)^2} \mathrm{tr}\left\{\mathbf{U}^{(m)}\right\} \right). \tag{S3.12}$$

On the other hand, $\alpha$ update takes a relatively simple form:

$$\alpha = \frac{2(L-1)M}{\left\langle \|\mathbf{C}_{2:L,:}\|^2 \right\rangle_{q(\mathbf{C}|M)}}. \tag{S3.13}$$

In brief, we start with an initial set of the hyperparameters $\boldsymbol{\theta}$ and $\alpha$, and keep alternating between variational posterior inference (E-step) and hyperparameter update (M-step) until the hyperparameters or the variational free energy stabilizes. Once the suitable hyperparameters are obtained, the variational inference is run for the last time providing us with the *final* oscillation components.

**Evaluation of variational free energy** Considering update rules of Eqs. S3.7, S3.13, S3.10 we can simplify the second term in the Eq S3.2 as:

$$\begin{aligned}\left\langle \log \frac{p\left(\mathbf{X}, \mathbf{Y}, \mathbf{C}, \mathbf{R}, \alpha, \boldsymbol{\theta} \mid M\right)}{q\left(\mathbf{X}, \mathbf{C}, \mathbf{R} \mid M\right)} \right\rangle_{q(\mathbf{X},\mathbf{C},\mathbf{R}|M)} &= -\frac{(L+2M)T}{2}\log 2\pi \\ &- \frac{T}{2}\log|\mathbf{Q}| - \frac{2M(L-1)}{2} - \left\langle \log q\left(\mathbf{X} \mid M\right)_{q(\mathbf{X}|M)} \right\rangle \\ &+ \frac{2M(L-1)}{2}\log\frac{\alpha}{2\pi} - \frac{2MT}{2} - \langle \log q(\mathbf{M} \mid m)\rangle \\ &+ \frac{\nu}{2}\log|\boldsymbol{\Psi}| + \gamma_{\nu,n} - \left(\frac{\nu+T}{2}\log|\boldsymbol{\Omega}| + \gamma_{\nu+T,n}\right).\end{aligned} \tag{S3.14}$$

After noting the following property of Gaussian random variable, $\mathbf{Z} \sim \mathcal{N}_p(\boldsymbol{\mu}_z, \boldsymbol{\Sigma}_z)$:

$$\langle \log p(\mathbf{Z}) \rangle_{p(\mathbf{Z})} = -\frac{1}{2}\log|(2\pi)\boldsymbol{\Sigma}_z| - \frac{p}{2}$$

Eq S3.14 can be further simplified as:

$$\begin{aligned}\left\langle \log \frac{p\left(\mathbf{X}, \mathbf{Y}, \mathbf{C}, \mathbf{R}, \alpha, \boldsymbol{\theta} \mid M\right)}{q\left(\mathbf{X}, \mathbf{C}, \mathbf{R} \mid M\right)} \right\rangle_{q(\mathbf{X},\mathbf{C},\mathbf{R}|M)} &= -\frac{LT}{2}\log 2\pi - \frac{T}{2}\log|\mathbf{Q}| + \frac{1}{2}\log|\boldsymbol{\Sigma}_x| \\ &+\frac{1}{2}\log|\alpha\boldsymbol{\Sigma}_c| + \frac{\nu}{2}\log|\boldsymbol{\Psi}| + \gamma_{\nu,n} - \left(\frac{\nu+T}{2}\log|\boldsymbol{\Omega}| + \gamma_{\nu+T,n}\right)\end{aligned} \tag{S3.15}$$

We note here that the log-determinant of $\boldsymbol{\Sigma}_x$ could be computed while solving Eq S3.8 exploiting the tri-diagonal structure of ${\boldsymbol{\Sigma}_x}^{-1}$. Similarly, the log-determinant of $\boldsymbol{\Sigma}_c$ could be computed while solving Eq S3.9 using the properties of Kronecker products. Matrices $\boldsymbol{\Phi}$ and $\boldsymbol{\Omega}$ are in the sensor space dimension, thus log-determinant computation is relatively cheaper. The remaining matrix $\mathbf{Q}$ is a diagonal matrix, thus its log-determinant can be computed easily. As a whole, the variational free energy for model structure $M$ can be computed as a by-product of the iterative updates.

**Initialization of parameters and hyper parameters** Because the ensemble log-likelihood (hence the negative variational free energy) is not concave in general, initial values for the parameters and hyperparameters must be carefully chosen. We use the interpretation of the MVAR processes as a superimposition of multiple oscillatory components [13–15].

First of all, we estimate $\boldsymbol{\Psi}$ using the fact that for single channel noisy recording, $y^{(l)}$ where $S_{y^{(l)}y^{(l)}}(f)$ denotes the power spectrum density (PSD) of the signal:

$$10 \log_{10} \frac{R}{f_s} = \lim_{f \to \infty} S_{y^{(l)}y^{(l)}}(f). \tag{S3.16}$$

We generalize S3.16 to multichannel recording by high-pass filtering the M/EEG recording at $f_s - \Delta f$, ($\Delta f > 0$), and estimating inverse-Wishart parameters from the sample covariance of the high-pass filtered data, $\widetilde{\mathbf{y}}_t$. We set $\nu = T$, and $\Psi = \sum_{t=1}^{T} \mathbf{x}_t \mathbf{x}_t^\top$. We also initialize the noise covariance matrix as $\mathbf{R} = \sum_{t=1}^{T} \mathbf{x}_t \mathbf{x}_t^\top / T$.

We then fit an MVAR model of order $p$ on the multichannel data. Order $p$ is chosen according to Akaike Information Criteria [16]. We perform eigendecomposition of the companion form of the MVAR parameter matrix [13] to yield $\boldsymbol{\lambda}$, $\mathbf{V}$ and $\mathbf{W}$ as eigenvalues, right and left eigenvectors. We choose the eigenvalues with $\Im\{\lambda^{(m)}\} > 0$, and corresponding right eigenvectors $\mathbf{w}^{(m)}$ and collect them as $\widehat{\boldsymbol{\lambda}}$ and $\widehat{\mathbf{W}}$ (assume there are $M$ such eigenvalues). The frequency and damping parameters are initialized as:

$$f^{(m)} = \arg\left(\lambda^{(m)}\right) \frac{f_s}{2\pi} \qquad\qquad a^{(m)} = |\lambda^{(m)}|.$$

For ${\sigma^{(1)}}^2, {\sigma^{(m)}}^2, \cdots, {\sigma^{(M)}}^2$, we use the linear combination of the theoretical PSD of the oscillatory components that approximates the mutitaper PSD [17] of the leading time-series the best. In short, we compute the mutitaper PSD of the leading time-series and the theoretical PSDs of the oscillatory components [18, Supplementary Information] on same frequency grid of spacing $f_s/(2K)$. The mutitaper PSD, $\boldsymbol{\rho}$ is a $1 \times K$ vector whereas the theoretical PSDs, $\boldsymbol{\phi}^{(m)}$ form a $M \times K$ matrix $\boldsymbol{\Phi}$. We initialize the ${\sigma^{(1)}}^2, {\sigma^{(m)}}^2, \cdots, {\sigma^{(M)}}^2$ as the solution of the following linear equation:

$$\begin{bmatrix} {\sigma^{(1)}}^2 & {\sigma^{(m)}}^2 & \dots & {\sigma^{(M)}}^2 \end{bmatrix} \boldsymbol{\Phi} = \boldsymbol{\rho}. \tag{S3.17}$$

In order to select a subset of the discovered $M$ oscillatory components, we sort the oscillatory components (and columns of the $\widehat{\mathbf{W}}$) according to their theoretical contribution (most to least):

$$\sigma^{2^{(m)}} \|\widehat{\mathbf{w}}^{(m)}/\widehat{w}_1^{(m)}\|_2 \sum_f \boldsymbol{\phi}^{(m)}(f), \tag{S3.18}$$

and select the leading oscillatory parameters (and columns of the $\widehat{\mathbf{W}}$).

With S3.17, the spatial mixing components for the leading time-series becomes $\mathbf{c}_{l,m} = \begin{bmatrix} 1 & 0 \end{bmatrix}$. The rest of the elements of spatial mixing matrix, $\mathbf{C}$, are initialized from $\widehat{\mathbf{W}}$ as:

$$\mathbf{c}_{l,m} = \left[ \Re\left(\frac{\widehat{\mathbf{w}}_l^{(m)}}{\widehat{\mathbf{w}}_1^{(m)}}\right) \quad -\Im\left(\frac{\widehat{w}_l^{(m)}}{\widehat{w}_1^{(m)}}\right) \right]$$

Lastly, $\alpha$ is initialized the $2(L-1)M/\|\mathbf{C}_{2:L}\|_{Fro}$ with the above-mentioned $\mathbf{C}$.

**Model structure posterior** Finally, it is easy to verify that the following discrete distribution maximizes Eq S3.2:

$$q(M) = \frac{p(M) \exp - \left\langle \log \frac{p(\mathbf{X},\mathbf{Y},\mathbf{C},\mathbf{R},\alpha,\boldsymbol{\theta}|M)}{q(\mathbf{X},\mathbf{C},\mathbf{R}|M)} \right\rangle_{q(\mathbf{X},\mathbf{C},\mathbf{R}|M)}}{\sum_{M'=1}^{M_{max}} p(M') \exp - \left\langle \log \frac{p(\mathbf{X},\mathbf{Y},\mathbf{C},\mathbf{R},\alpha,\boldsymbol{\theta}|M')}{q(\mathbf{X},\mathbf{C},\mathbf{R}|M')} \right\rangle_{q(\mathbf{X},\mathbf{C},\mathbf{R}|M')}}, \tag{S3.19}$$

which can be computed readily once the final variational inferences are made for $M = 1, \cdots, M_{\max}$. The negative variational free-energy of the modelling framework is given by,

$$\mathcal{F} = \sum_{M'=1}^{M_{max}} p(M') \exp - \left\langle \log \frac{p\left(\mathbf{X}, \mathbf{Y}, \mathbf{C}, \mathbf{R}, \alpha, \boldsymbol{\theta} \mid M'\right)}{q\left(\mathbf{X}, \mathbf{C}, \mathbf{R} \mid M'\right)} \right\rangle_{q(\mathbf{X},\mathbf{C},\mathbf{R}|M')}. \tag{S3.20}$$

**Empirical Bayes inference and model selection** Since the model order is not known, we estimate the model order, $M$ by maximizing the approximate posterior distribution of model order, $q(M)$, and treat $M$ as if it were known to be equal to this estimate. The same procedure is followed for the hyperparameters, for which we set them to the values that maximizes the negative variational free energy. This way of determining model order or estimating unknown parameters comes under the umbrella of empirical Bayes framework.[19–22] Briefly, it provides a first order approximation to the posterior mean of these quantites, and neglects the uncertainity (i.e. second order statistics) of the estimated quantities.

##### Comparison of OCA to traditional approaches in experimental EEG data

We compare OCA with an oscillation finding approach based on channel-wise power spectral density visualization and ICA in a subset of real human EEG recording, taken from Fig 4 to provide empirical justification for OCA. For Fig S4.1, we used a 3.5 s long EEG segment during the maintenance of 2 µg effect-site concentration of propofol. We expect strong frontal alpha activity and overall increase in slow activity as per existing literature [23, 24].

The frequency domain approach examined here is based on the popular multitaper method [17]: given a bandwidth parameter, power spectrum density is computed for individual channels and a butterfly plot is made (see Fig 3A left panels). The oscillatory alpha activities are then identified visually (red overlay) as the peaks in these power spectrum plots, and their scalp distribution are obtained by averaging channel-wise power within given frequency bands around the peaks as shown in Fig 3A right panels. We chose three different bandwidth, 1 Hz, 2 Hz and 5 Hz to demonstrate the inconsistency of frequency domain methods in 'producing' peaks in the line plots of the spectrum. Spectrum with 1 Hz bandwidth (top row) exhibits a lot of spurious peaks, whereas spectrum with 5 Hz bandwidth (bottom row) exhibits wide peaks, possibly encompassing multiple peaks. This inconsistency of ad-hoc visual identification of peaks makes it subjective. Similarly, the topographic plots in Fig 3A demonstrates similar inconsistency: as estimation bandwidth increases (i.e., spectral resolution decreases), the topographic maps becomes less susceptible to spurious power leakage until the estimation bandwidth matches the bandwidth of underlying processes (compare middle row to top row). As the estimation bandwidth further increases, a severe spectral leakage renders the topographic maps largely uninformative of underlying separable oscillatory sources (bottom row).

ICA decomposition on this multichannel recordings identifies components that are neither slow oscillation nor alpha oscillation but combination of multiple oscillations, as one would expect based on the observation that cumulative histograms of EEG recordings is approximately Gaussian [25]. We use 'extended infomax' [26] implementation provided by MNE-python 1.2 [27], the leading 4 ICA components are shown in Fig S4.1B.

Lastly, OCA is able to identify several oscillatory components (in both slow and alpha band) from this multichannel data, extract their individual time-series and topographic distribution. OCA is able to do so due to its explicit modelling of temporal dynamics in form of the structured state-space representation. Fig S4.1B shows leading 4 components whose estimated center frequencies are within alpha band (9 Hz to 13 Hz). This also demonstrates that OCA is bandwidth-free, i.e., it adjusts estimation bandwidth to match the bandwidths of the underlying oscillations in a data-driven way as evident from the spectrum plots of the estimated oscillation time-courses.

#### References


1. Galinsky VL, Frank LR. Universal Theory of Brain Waves: From Linear Loops to Nonlinear Synchronized Spiking and Collective Brain Rhythms. Physical Review Research. 2020;2(2):023061. doi:10.1103/PhysRevResearch.2.023061.

2. Rosenblum MG, Pikovsky AS, Kurths J. Phase synchronization in driven and coupled chaotic oscillators. IEEE Transactions on Circuits and Systems I: Fundamental Theory and Applications. 1997;44(10):874–881. doi:10.1109/81.633876.

3. Muirhead RJ. Aspects of Multivariate Statistical Theory. Wiley Series in Probability and Mathematical Statistics. New York: Wiley; 1982.

4. Neal RM, Hinton GE. A View of the EM Algorithm That Justifies Incremental, Sparse, and Other Variants. In: Jordan MI, editor. Learning in Graphical Models. Dordrecht: Springer Netherlands; 1998. p. 355–368.

5. Quinn A, Šmídl V. The Variational Bayes Method in Signal Processing. Signals and Communication Technology Ser. New York Boulder: Springer NetLibrary, Inc.; 2006.

6. Attias H. Inferring Parameters and Structure of Latent Variable Models by Variational Bayes. In: Proceedings of the Fifteenth Conference on Uncertainity in Artificial Intelligence; 1999. p. 21–30.

7. Fahrmeir L. Posterior Mode Estimation by Extended Kalman Filtering for Multivariate Dynamic Generalized Linear Models. Journal of the American Statistical Association. 1992;87(418):501–509. doi:10.1080/01621459.1992.10475232.

8. Boyd SP, Vandenberghe L. Convex Optimization. Cambridge, UK ; New York: Cambridge University Press; 2004.

9. Asif A, Moura JMF. Inversion of Block Matrices with Block Banded Inverses: Application to Kalman-Bucy Filtering. In: 2000 IEEE International Conference on Acoustics, Speech, and Signal Processing. Proceedings (Cat. No.00CH37100). vol. 1. Istanbul, Turkey: IEEE; 2000. p. 608–611.

10. Jain J, Li H, Cauley S, Koh CK, Balakrishnan V. Numerically Stable Algorithms for Inversion of Block Tridiagonal and Banded Matrices. Department of Electrical and Computer Engineering Technical Reports. 2007;.

11. Dempster AP, Laird NM, Rubin DB. Maximum Likelihood from Incomplete Data Via the *EM* Algorithm. Journal of the Royal Statistical Society: Series B (Methodological). 1977;39(1):1–22. doi:10.1111/j.2517-6161.1977.tb01600.x.

12. Shumway RH, Stoffer DS. An Approach To Time Series Smoothing And Forecasting Using The Em Algorithm. Journal of Time Series Analysis. 1982;3(4):253–264. doi:10.1111/j.1467-9892.1982.tb00349.x.

13. Neumaier A, Schneider T. Estimation of Parameters and Eigenmodes of Multivariate Autoregressive Models. ACM Transactions on Mathematical Software. 2001;27(1):27–57. doi:10.1145/382043.382304.

14. Matsuda T, Komaki F. Multivariate Time Series Decomposition into Oscillation Components. Neural Computation. 2017;29(8):2055–2075. doi:10.1162/neco_a_00981.

15. Quinn AJ, Green GGR, Hymers M. Delineating Between-Subject Heterogeneity in Alpha Networks with Spatio-Spectral Eigenmodes. NeuroImage. 2021;240:118330. doi:10.1016/j.neuroimage.2021.118330.

16. Cavanaugh JE, Neath AA. The Akaike Information Criterion: Background, Derivation, Properties, Application, Interpretation, and Refinements. WIREs Computational Statistics. 2019;11(3). doi:10.1002/wics.1460.

17. Babadi B, Brown EN. A Review of Multitaper Spectral Analysis. IEEE Transactions on Biomedical Engineering. 2014;61(5):1555–1564. doi:10.1109/TBME.2014.2311996.

18. Soulat H, Stephen EP, Beck AM, Purdon PL. State Space Methods for Phase Amplitude Coupling Analysis. Scientific Reports. 2022;12(1):15940. doi:10.1038/s41598-022-18475-3.

19. Robbins H. The Empirical Bayes Approach to Statistical Decision Problems. The Annals of Mathematical Statistics. 1964;35(1):1–20.

20. Efron B, Morris C. Stein's Estimation Rule and its Competitors—An Empirical Bayes Approach. Journal of the American Statistical Association. 1973;68(341):117–130. doi:10.1080/01621459.1973.10481350.

21. Efron B, Morris C. Data Analysis Using Stein's Estimator and its Generalizations. Journal of the American Statistical Association. 1975;70(350):311–319. doi:10.1080/01621459.1975.10479864.

22. Morris CN. Parametric Empirical Bayes Inference: Theory and Applications. Journal of the American Statistical Association. 1983;78(381):47–55. doi:10.1080/01621459.1983.10477920.

23. Cimenser A, Purdon PL, Pierce ET, Walsh JL, Salazar-Gomez AF, Harrell PG, et al. Tracking Brain States under General Anesthesia by Using Global Coherence Analysis. Proceedings of the National Academy of Sciences. 2011;108(21):8832–8837. doi:10.1073/pnas.1017041108.

24. Purdon PL, Pierce ET, Mukamel EA, Prerau MJ, Walsh JL, Wong KF, et al. Electroencephalogram Signatures of Loss and Recovery of Consciousness from Propofol. Proc Natl Acad Sci U S A. 2013;110(12):E1142–51. doi:10.1073/pnas.1221180110.

25. Brookes MJ, Woolrich M, Luckhoo H, Price D, Hale JR, Stephenson MC, et al. Investigating the Electrophysiological Basis of Resting State Networks Using Magnetoencephalography. Proceedings of the National Academy of Sciences. 2011;108(40):16783–16788. doi:10.1073/pnas.1112685108.

26. Lee TW, Girolami M, Sejnowski TJ. Independent Component Analysis Using an Extended Infomax Algorithm for Mixed Subgaussian and Supergaussian Sources. Neural Computation. 1999;11(2):417–441. doi:10.1162/089976699300016719.

27. Gramfort A, Luessi M, Larson E, Engemann DA, Strohmeier D, Brodbeck C, et al. MNE Software for Processing MEG and EEG Data. NeuroImage. 2014;86:446–460. doi:10.1016/j.neuroimage.2013.10.027.

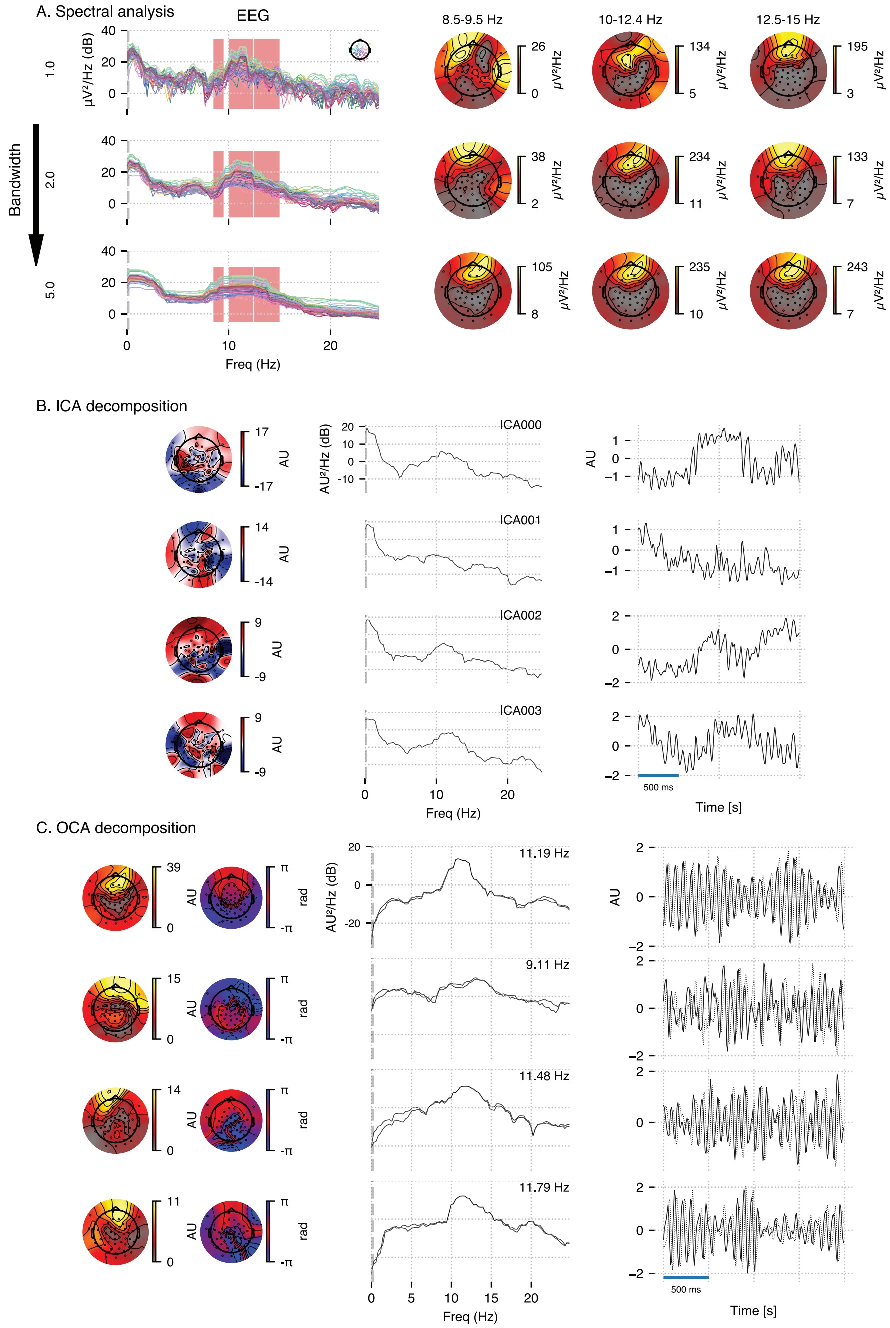


**Figure S4.1.** Empirical justification for OCA in analyzing real data. A 3.5 s long EEG recording during propofol-induced anesthesia (effect-site concentration of 2 μg) is considered for this demonstration. **A.** power spectral density using multitaper method (for varying time-bandwidth product) of the EEG recording, and power distribution over the EEG sensors in the marked frequency bands (red overlay) around visually identifiable peaks; **B.** Four leading ICA components (left, middle and right columns show topographic maps, power spectrum density and time-courses respectively); **C.** Four leading OCA components within alpha band (the topographic maps show the magnitude (left) and phase (right), while line plots show power spectrum density (left) and time-courses (right) respectively.)
